# MrtR of *Mesorhizobium tianshanense* reveals both activation and inhibition mechanisms of a LuxR-type quorum sensing receptor

**DOI:** 10.64898/2026.05.28.728602

**Authors:** Irene M. Stoutland, Helen E. Blackwell

**Affiliations:** Department of Chemistry, University of Wisconsin–Madison, 1101 University Avenue, Madison, WI 53706, USA

**Keywords:** *N*-acyl L-homoserine lactone, autoinducer, cell-cell signaling, quorum sensing, rhizobia, transcription factor

## Abstract

Quorum sensing (QS) enables common gram-negative bacteria to coordinate collective behaviors through small molecule signals, yet how these signals tune receptor activity remains incompletely understood. Here, we define a mechanism by which ligand structure controls function in a LuxR-type QS receptor. Using structural and biochemical analyses, we investigate MrtR from *Mesorhizobium tianshanense* and show that ligand acyl-chain length governs receptor assembly and activity. We present full-length structures of MrtR bound to activating and inhibitory ligands, revealing a switch in oligomeric state. Long-chain (C14) *N*-acyl L-homoserine lactones (AHLs) act as agonists by promoting intra- and inter-subunit interactions that lead to homodimerization and DNA binding. In contrast, shorter (C8) AHLs fail to promote these contacts, favoring a monomeric, inactive state. Ligands of intermediate length produce graded responses consistent with partial dimer stabilization. Biochemical measurements of DNA binding, thermostability, and oligomerization, together with targeted mutagenesis, support this model and establish the functional importance of key structural contacts. These findings provide the first structural comparison of a full-length LuxR-type receptor bound to both agonist and antagonist. Our findings expand the known structural and mechanistic diversity of the LuxR family and suggest mechanistic similarities between structurally distinct receptors.

**SIGNIFICANCE:** Quorum sensing (QS) regulates diverse bacterial behaviors, and LuxR-type receptors are attractive targets for applications ranging from antivirulence strategies to synthetic biology and agriculture. Despite intense interest in developing chemical modulators of these systems, the molecular basis by which small molecules agonize or antagonize LuxR-type receptors remains poorly understood. Here, we investigate the LuxR-type receptor MrtR and report crystal structures of the full-length receptor bound to an agonist and an antagonist, revealing how structurally similar compounds produce opposing outcomes. Notably, MrtR exhibits an unprecedented dimerization interface mediated by a ligand-responsive loop that undergoes large conformational changes. These findings establish a new structural framework for understanding signal discrimination in LuxR-type receptors and may enable rational reprogramming of QS in natural and engineered systems.

## INTRODUCTION

Quorum sensing (QS) is a system of cell-cell communication used by many common bacterial species to regulate cell density-dependent phenotypes (1, 2). LuxI/R QS involves a LuxI-type synthase that produces an *N*-acyl L-homoserine lactone (AHL) signaling molecule and an intracellular LuxR-type receptor that regulates gene expression in response to AHL concentration (3). Diverse species of gram-negative bacteria use LuxI/R QS to regulate phenotypes such as bioluminescence, biofilm formation, virulence, natural product synthesis, and symbiosis (3, 4). Because LuxI/R systems are targets for applications in antivirulence (5), antibiofouling (4), and biotechnology (6), there is substantial interest in developing synthetic agonists and antagonists to control LuxR-type receptor activity (4, 5, 7-12).

Despite widespread attention to LuxI/R systems, a lack of understanding about the molecular mechanisms by which ligand binding controls LuxR activity presents a barrier to improved strategies to modulate LuxI/R QS. The poor solubility of LuxR-type receptors in vitro, especially in the absence of AHL ligand, has hampered previous studies of LuxR mechanism, and structural work has often required domain truncations and/or stabilizing ligands (13, 14). Of the thousands of putative LuxR-type receptors (15), nine have structures available in the PDB, many comprising a truncated ligand-binding domain (LBD). Only CviR of *Chromobacterium subtsugae* and CviR’ of *C. violaceum* have been crystallized in the presence of antagonists, and their corresponding agonist-bound structures include only the LBD (13). A comparison of full-length structures of a LuxR-type receptor in the presence of agonist vs. antagonist could illuminate the mechanisms of small molecule activation and inhibition, but no such structures are currently available.

To address this challenge, we sought to find a LuxR homolog that could facilitate in vitro and structural work. Thousands of putative LuxR-type receptor sequences identified using BLAST (16) were screened using Crysalis (17) to identify promising candidates for structural work. One of the top hits was MrtR from *Mesorhizobium tianshanense*, a nitrogen-fixing rhizobia that requires MrtR for the formation of symbiotic nodules on *Glycyrrhiza uralensis* (licorice) (18). Proteins 95-100% identical to MrtR are present in various species of rhizobia (**Table S1**) that use QS to regulate plasmid transfer (19), growth rate (20), and symbiotic nitrogen fixation (21, 22). Previous work by Zhu and co-workers has shown that MrtR is soluble in the absence of ligand but requires AHL for activity (23), giving us confidence that MrtR could facilitate in vitro mechanistic studies.

Herein we report the biochemical and structural characterization of MrtR’s response to a set of AHL ligands. We identify AHLs with a range of effects on MrtR’s transcriptional activity, measure how these AHLs affect DNA binding and thermal stability, show that antagonists exert their effects by inhibiting MrtR dimerization, and present structures of full-length MrtR bound to both agonists and antagonists. The structures support a mechanistic model in which binding of long-acyl chain AHL agonists, but not short-acyl chain antagonists, stabilizes residues at MrtR’s dimerization interface, inducing dimerization and receptor activation.

## RESULTS

### *E. coli* reporter screen reveals strong agonists and antagonists of MrtR

Naturally occurring AHLs share an L-homoserine lactone (HSL) head group and vary in the length and structure of the acyl tail (24). LuxR homologs typically respond strongly to their native AHL, produced by their cognate LuxI-type synthase (3, 4). MrtR activates expression of its cognate synthase gene, *mrtI*, in response to *N*-(3*R*-hydroxy)-7-*cis*-tetradecenoyl L-homoserine lactone (7*Z*-3*R*-OHC14) (25). We used a previously described (25) MrtR reporter in *E. coli* to test a panel of AHLs for their ability to activate or inhibit MrtR transcriptional activation of the *mrtI* promoter (see Methods). DNA sequences identical or highly similar to this promoter sequence are present in various species of rhizobia (**Table S2**), supporting the relevance of this reporter system to biological interactions in multiple species.

Naturally occurring 14-carbon AHLs acted as agonists of MrtR, and 8-carbon AHLs acted as antagonists of MrtR when competed against 7*Z*-3OHC14 (mix of 3*R* and 3*S* diastereomers) (**Figure 1A** and **B**). AHLs with tail lengths between 8 and 14 carbons showed intermediate activity that varied with oxidation state; notably, both 3*R* and 3*S* diastereomers of 3OHC12 were highly activating, while C12-HSL and 3OC12 acted as partial agonists, showing both agonism and antagonism activity. A panel of synthetic AHLs from our in-house AHL analog libraries was also screened for activity (**Figure S1, Table S3**), and several showed moderate antagonism, while the chloro aryl AHL analog H23 strongly antagonized activity (**Figure 1A** and **B**). Compounds with strong agonism or antagonism activity were further examined for dose-dependent effects on activity in the *E. coli* reporter (**Table 1, Figures S2** and **S10**). The most potent agonists were 7*Z*-3OHC14 and 3OHC14, with ∼2 nM EC_50_ values, while 3OC14 and C14 produced EC_50_ values of ∼50-100 nM. The stereochemistry of the 3-OH group affected potency of 3OHC12: the 3*R* diastereomer was almost 9-fold more potent than the 3*S* diastereomer, as previously observed in AbaR (26). The three antagonists tested, 3OC8, 3OHC8, and H23, produced comparable IC_50_ values (∼700 nM) and full inhibition when competed against 8 nM 7*Z*-3OHC14, representing some of the most potent and efficacious AHL-type inhibitors uncovered for a LuxR-type receptor (12).

**Table 1.**
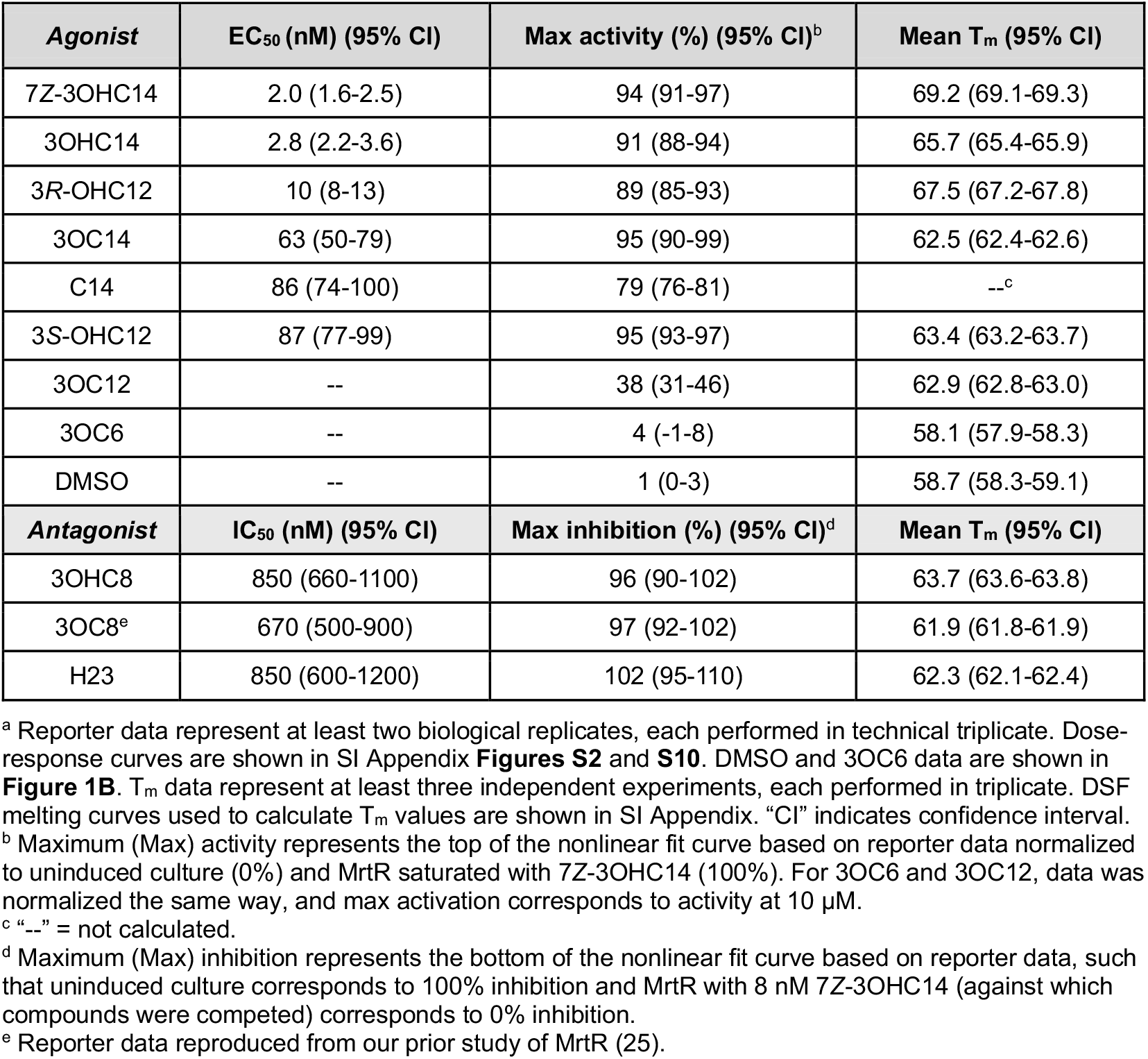
EC_50_ and IC_50_ values and efficacies in cells and melting temperatures (T_m_) in vitro for select agonists and antagonists at MrtR.^a^.

**Figure 1.**
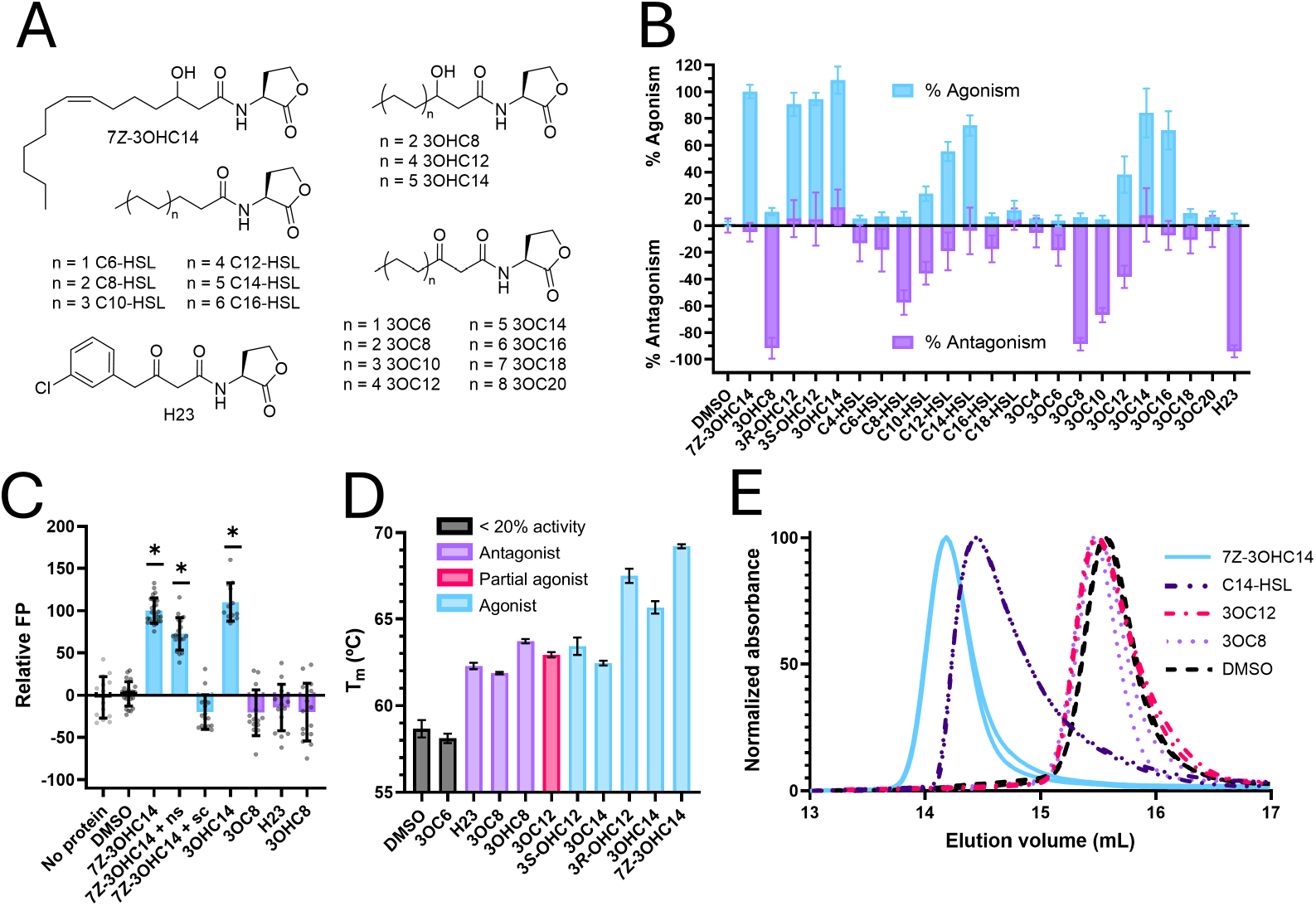
Activity of MrtR in cells and properties in vitro. **(A)** A subset of the compounds assayed for activity in MrtR assays. Additional compounds and activity data are shown in **Figure S1**; compound citations are listed in **Table S3. (B)** Agonism and antagonism activity of compounds in an *E. coli* MrtR transcriptional reporter. Percent agonism represents transcription of *mrtI*:GFP with each compound at 10 μM, normalized to uninduced culture (0%) and MrtR saturated with 7*Z*-3OHC14 (100%). Percent antagonism represents transcription of *mrtI*:GFP with 10 μM of each compound competed against 8 nM 7*Z*-3OHC14, normalized to uninduced culture (-100%) and 8 nM 7*Z*-3OHC14 (0%). Data represent at least two biological replicates, each performed in technical triplicate, except for H23 agonism, which represents one biological replicate. **(C)** Fluorescence polarization (FP) data measuring MrtR (400 nM) binding to the FAM-labeled *mrtI* promoter (5 nM) with 100 μM compound, normalized to FP in the presence of DMSO (0%) and FP in the presence of 7*Z*-3OHC14 (100%). Antagonists are shown in purple, agonists in blue, and conditions without compound in gray. A section of the pUC18 multiple cloning site and unlabeled *mrtI* promoter were used for nonspecific (ns) and specific competitor (sc) controls, respectively, and both were included at 500 nM in the indicated samples. Asterisks indicate that relative FP is significantly different from the DMSO condition based on an ANOVA with p ≤ 0.01. Data represent at least three experiments of at least three replicates. **(D)** Differential scanning fluorimetry (DSF) data for MrtR at 5 μM in the presence of 100 μM compound. Agonists (blue) are ordered from left to right based on their potency (low to high) in a transcriptional reporter assay (**Table 1**). Data represent at least three experiments, each performed in triplicate. T_m_ values listed in **Table 1. (E)** Size exclusion chromatograms of MrtR in the presence of indicated compounds. Both duplicate runs are shown. Traces are normalized to the smallest (0) and largest (100) values within the range shown. Full traces are shown in SI Appendix and calculated molecular weight values in **Table S4. (All bar graphs)** Error bars indicate standard deviation.

### Agonists but not antagonists promote MrtR:DNA binding, and weak agonists and antagonists induce a similar degree of MrtR thermostability

To begin to understand why certain compounds act as agonists and others as antagonists, we measured the effect of compounds on MrtR:DNA binding and MrtR thermostability in vitro. MrtR was expressed with an N-terminal His6-SUMO tag and purified by immobilized metal affinity chromatography (IMAC). The tag was cleaved using SUMO protease Ulp1 and removed with another round of IMAC, resulting in pure MrtR with the native N-terminus (see SI Appendix). Apo-MrtR was soluble up to at least 7 mg/mL, allowing us to purify the apo form and incubate with compounds of interest for in vitro assays.

A fluorescence polarization (FP) assay using a 42 base pair probe corresponding to the MrtR binding site on the *mrtI* promoter (23) was used to examine MrtR:DNA binding in the presence of agonists and antagonists. As expected from the reporter data, agonists 7*Z*-3OHC14 and 3OHC14 promoted DNA binding, while antagonists 3OC8, 3OHC8, and H23 did not (**Figure 1C**). Selected compounds were also tested for their ability to increase MrtR thermostability (**Figure 1D** and **Table 1**). The most potent agonists, 7*Z*-3OHC14, 3OHC14, and 3*R-*OHC12 were the most thermostabilizing, consistent with high binding affinity for these compounds. T_m_ values did not differ substantially between weak agonists (3*S*-OHC12 and 3OC14), a partial agonist (3OC12), and antagonists (3OC8, 3OHC8, and H23), indicating that the ability of these compounds to thermostabilize MrtR does not determine whether they act as agonists or antagonists.

### Agonists but not antagonists induce MrtR dimerization

LuxR-type receptors typically bind to DNA as homodimers (15), and agonist binding has previously been shown to induce MrtR dimerization (23). Size exclusion chromatography (SEC) was used to estimate the molecular weight of MrtR complexes in the presence of different compounds (**Figure 1E** and **Table S4**). MrtR has a molecular weight of 27.2 kDa based on sequence and eluted as a monomer in the absence of compound (∼26 kDa) and in the presence of antagonist 3OC8 (∼27 kDa). In contrast, in the presence of agonist 7*Z*-3OHC14, MrtR eluted as a dimer (∼48 kDa). As 48 kDa is slightly lower than the expected 54 kDa for an MrtR dimer, agonist-bound MrtR may adopt a highly compact dimer conformation or exist in a dynamic equilibrium, as has been observed for other LuxR-type receptors (27).

Partial agonist 3OC12, which shows ∼38% agonism in the cell-based reporter (**Table 1**), caused MrtR to elute as a monomer (∼27 kDa). Dimerization can be sensitive to protein concentration and environmental factors, and although 3OC12 is able to activate MrtR in cells, it was unable to induce dimerization under our in vitro conditions. With partial agonist C14-HSL (∼79% agonism; **Table 1**), MrtR eluted at ∼43 kDa and showed substantial tailing, which suggests a monomer-dimer equilibrium with a slower association rate or faster dissociation rate than MrtR:7*Z*-3OHC14 (28) and is congruent with the partial agonism activity of C14-HSL in the reporter.

In vitro chemical crosslinking experiments also supported that agonists and antagonists differ in their ability to induce MrtR dimerization. MrtR bound to various ligands was subjected to BS(PEG)_5_ and crosslinking was measured by SDS-PAGE. Crosslinks were observed in the presence of agonists 7*Z*-3OHC14, 3OHC14, 3*R-*OHC12, and 3*S-*OHC12 but not in the absence of compound (DMSO) or in the presence of antagonists 3OC8 or 3OHC8 or partial agonist 3OC12 (**Figure S3**). Taken together, these crosslinking and SEC data strongly suggest that the ability of a compound to induce MrtR dimerization determines its degree of agonism activity and differentiates agonists from antagonists.

### Crystal structures inform MrtR mechanism of ligand response

We next turned to structural analysis to gain insight into the molecular mechanisms by which agonists and antagonists act on MrtR. Our initial forays involved the use of AlphaFold 3 (29), which produced an MrtR homodimer model with unusually high confidence compared to other LuxR-type receptors that we have modeled (**Figure S4**) yet with a LBD dimerization interface that differed starkly from existing LuxR structures. Remarkably, the predicted dimerization interface matched our experimentally determined agonist-bound MrtR X-ray crystal structures (**Figure S4**) and the full-length monomer or separate domains from the AlphaFold model were used for molecular replacement for both agonist- and antagonist-bound structures. We solved the crystal structures of MrtR bound to agonists 7*Z*-3OHC14 (at 3.06 Å) and 3OHC14 (at 3.36 Å), which were highly similar (**Figure S5**), and structures of MrtR bound to antagonists 3OHC8 (at 1.18 Å) and 3OC8 (at 1.44 Å). One of the datasets with 3OC8 (at 1.22 Å) showed additional density in the binding site, likely corresponding to a tetraethylene glycol (TEG) molecule (data collection and refinement statistics in **Table S5**).

Overall, MrtR shows the same helix-turn-helix DNA-binding domain (DBD) and Per-ARNT-Sim ligand-binding domain (LBD) observed in reported structures of other LuxR-type receptors (13, 30-32). As expected based on the SEC and crosslinking data, MrtR crystallized as monomer in the presence of antagonists and as a dimer in the presence of agonists according to PDBePISA (33) analysis (**Tables S6** and **S7**). In agonist-bound MrtR, both the LBD and DBD are dimerized, and the MrtR LBD dimerization interface is distinct relative to available LuxR-type receptor structures (discussed further below), while the DBD dimerization interface is highly similar (**Figures S6** and **S7**). In contrast to agonist-bound MrtR, antagonist-bound MrtR shows the DBD buried against the LBD (**Figure 2A**), occluding portions of both the fourth helix of the DBD, which engages in dimerization in agonist-bound MrtR, and the third helix of the DBD, which engages in DNA binding in reported structures of DNA-bound LuxR-type receptors (i.e., TraR and RhlR, **Figure S6**) (34, 35). Crystals of apo-MrtR produced low-quality diffraction data, and further structure refinement was not pursued, but preliminary analysis indicates a different space group than antagonist-bound structures and shows that apo-MrtR adopts an overall domain orientation similar to antagonist-bound MrtR, implying that the observed LBD-DBD contacts are not an artifact of crystal packing.

**Figure 2.**
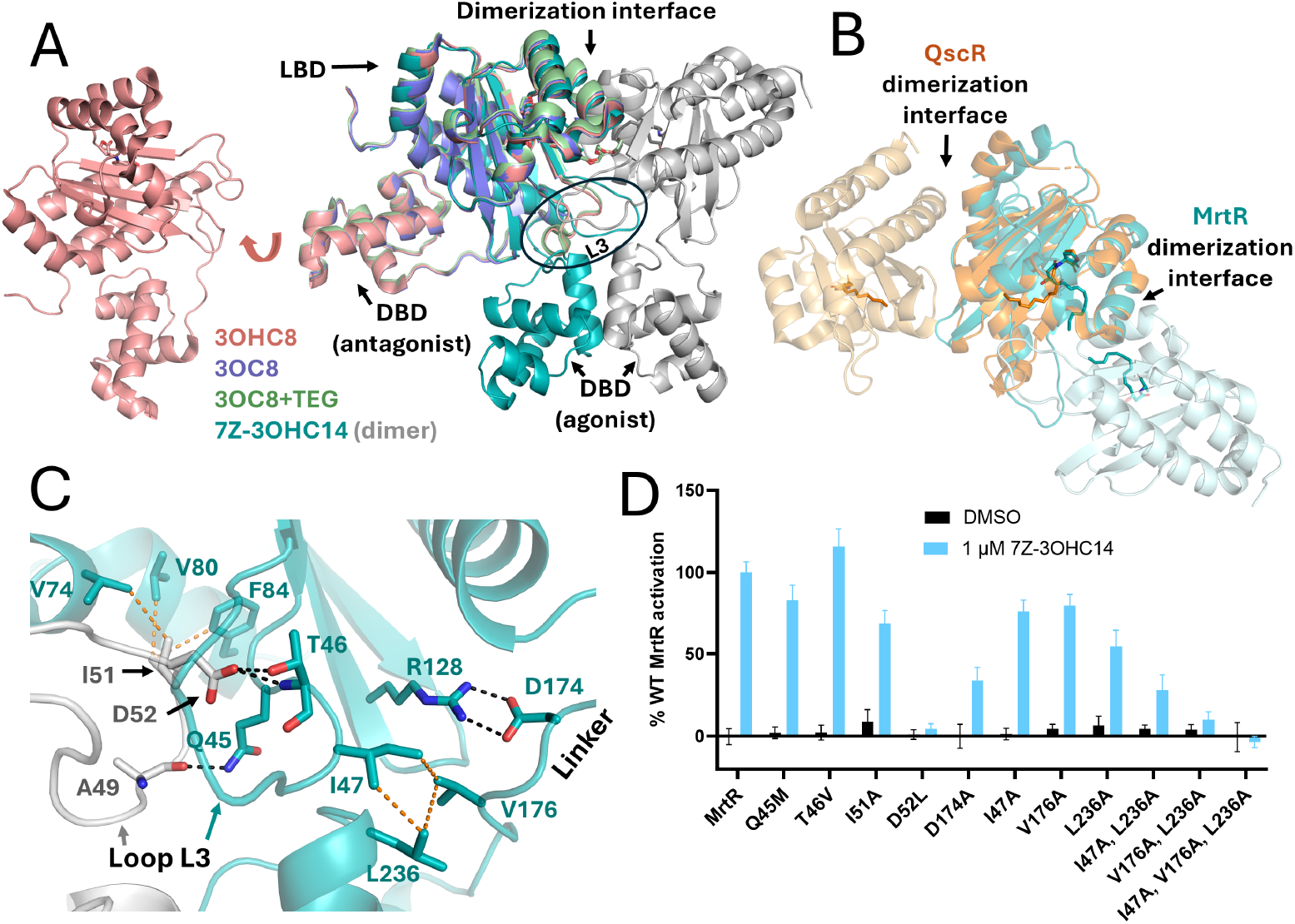
MrtR structure overview and loop L3 dimerization contacts. **(A)** Comparison of MrtR structures bound to 7*Z*-3OHC14 (teal), 3OHC8 (salmon), 3OC8 (violet), and 3OC8+TEG (green). Note differences in DBD orientation and L3 conformation. **(B)** Comparison of the MrtR (teal) and QscR (PBD ID: 3SZT (36), orange) LBD. Opposite monomers for each receptor are shown in a lighter shade of the same color. Despite distinct dimerization interfaces and ligand conformations, the ligand tail approaches the dimerization interface in both receptors. **(C)** MrtR:7*Z*-3OHC14 structure showing dimerization, linker, and inter-domain contacts probed by mutagenesis. Monomers are shown in teal and gray, as in panel **(A).** Hydrogen bonds and salt bridges (< 3 Å) are shown in black, and hydrophobic interactions are shown in orange (≤ 4.3 Å). **(D)** Transcriptional activation by point mutants of residues shown in panel **(C)** in an *E. coli* MrtR reporter. Data are normalized to uninduced culture (0%) and WT MrtR saturated with 7*Z*-3OHC14 (100%) and represent at least three biological replicates, each performed in technical triplicate. Error bars indicate standard deviation. T46V showed slightly higher maximum activity than WT MrtR, and all other mutants had maximum activity significantly lower than WT MrtR based on an ANOVA with Dunnett’s correction for multiple comparisons and p ≤ 0.01.

### The dimerization interface of MrtR differs from existing LuxR structures

The LBD dimerization interfaces among structurally characterized LuxR-type receptors differ only slightly and tend to involve helices α1 (res. ∼5-20) and/or α6 (res.∼145-165), as well as several loops (14). In contrast, the MrtR:7Z-3OHC14 dimerization interface involves loop L3 (between strands β1 and β2; res. ∼44-54), strand β2 (res. ∼55-58), and helix α3 (res. ∼62-71), which are located on the *opposite* side of the LBD from helices α1 and α6 (**Figures 2B** and **S7**). Interestingly, L3 plays an important role in other LuxR-type receptors as well, adopting different conformations based on bound ligand in LasR (37) and engaging in dimerization and LBD-DBD contacts in QscR (36).

In MrtR, L3 mediates intermonomer and interdomain contacts and shows a large conformational shift between agonist- and antagonist-bound structures. In MrtR:7*Z*-3OHC14, L3 adopts an extended position that is markedly different from other LuxR structures (**Figure S6**) and enables extensive dimerization contacts. In contrast, in MrtR:3OC8+TEG, L3 adopts a compact, partially helical conformation (**Figures 2A** and **S6**). The electron density for L3 in MrtR:3OHC8 suggests multiple conformations, and L3 was not modeled in MrtR:3OC8 due to insufficient density, suggesting conformational disorder in this region (**Figure S8**).

To verify the biological relevance of interactions observed in the MrtR crystal structures, conservative mutations were made to several residues in L3 that make dimerization contacts in the MrtR:7*Z*-3OHC14 crystal structure, and mutant activity was tested in an *E. coli* transcriptional reporter in the presence of 7*Z*-3OHC14. A D52L mutation caused the largest (96%) decrease in activity, while Q45M and I51A caused intermediate decreases in activity (17% and 32%, respectively), and T46V did not decrease activity, likely because D52 can hydrogen bond to either the side chain or backbone of T46 (**Figure 2C** and **2D**).

L3 also mediates intramonomer interactions between domains, specifically I47 of L3 in the LBD, V176 of the linker, and L236 of the DBD, in MrtR bound to agonist but not antagonist (**Figure 2C**). Mutation of each of these residues alone caused a minor to moderate decrease in activity (20-45%), while double mutants showed a larger reduction in activity (70-90%) and a triple mutant was inactive (**Figure 2D**). In addition, mutation of linker residue D174, which is positioned to hydrogen bond to R128 of the LBD, decreased activity by 66%. Together, these mutagenesis results indicate that specific interactions between the LBD, DBD, and linker are important for MrtR activation.

### Ligand binding in MrtR structures

Despite MrtR’s unique structural features, the headgroups of the bound AHLs make contacts that are conserved in other crystal structures of LuxR-type receptors: hydrogen bonds from the HSL carbonyl, amine nitrogen, and amide oxygen to W64, D77, and Y60, respectively. The tails of 7*Z*-3OHC14 and 3OHC14 both adopt a distinctly bent conformation when bound to MrtR (**Figure S5**), and the *cis*-alkene in the 7*Z*-3OHC14 tail may promote this kinked conformation.

The acyl tails of the AHLs bound to MrtR extend approximately perpendicular to the strands of the central β-sheet, similar to the AHL tail conformations in CviR (13), TraR (34), and SdiA (31) structures and unlike the 12-carbon AHLs bound to LasR (14) and QscR (36) (**Figures 2B** and **S9**). But while the ligand tail in CviR, TraR, and SdiA extends towards the solvent, the unique dimerization interface of MrtR means that the ligand tail extends towards the dimerization interface, even making contacts with the opposite monomer. Although the AHL tails in MrtR and QscR extend in different directions, the difference in dimerization interface between these receptors means that the AHL tails extend towards the dimerization interface in both cases and likely stabilize residues involved in dimerization (**Figure 2B**) (36, 38).

MrtR is the first LuxR-type receptor crystallized with a 3-hydroxy-substituted AHL (as opposed to 3-oxo or unsubstituted AHLs), and comparison of antagonist-bound MrtR structures reveals structural determinants of MrtR ligand preference. MrtR antagonists 3OHC8 and 3OC8 differ in hydrogen bonding contacts between the 3O or 3OH substituent and H42, suggesting a role for this residue in ligand specificity. Indeed, while MrtR is more sensitive to 3OH over 3O AHLs in an *E. coli* reporter assay, an H42L mutation reversed this preference (**Figure S10**). As H42L showed lower maximum activity than WT MrtR, H42 may also play a role in agonism by influencing the conformation of adjacent loop L3 (**Figures S11** and **S12**).

### MrtR agonists engage in more extensive contacts near the dimerization interface than antagonists

As introduced above, helix α3, strand β2, and loop L3 are important dimerization elements, and the longer tail of agonist 7*Z*-3OHC14 allows it to interact with these elements in ways that the shorter antagonists 3OHC8 and 3OC8 cannot (overlay shown in **Figure 3A**). Agonist 7*Z*-3OHC14 and antagonist 3OHC8 engage in similar interactions with MrtR through about the eighth carbon of the ligand tail, but thereafter, the longer tail of 7*Z*-3OHC14 engages in inter- and intramonomer contacts to P54 (L3) and L69 (α3). Interactions with P54 may influence the conformation of L3 to promote inter-monomer interactions by residues within L3. Likewise, contacts to residues along α3 and β2 may help position other residues along these secondary structural elements, such as R57, D62, V65, S66, L69, and L70, for favorable interactions with the opposite monomer (key contacts highlighted in **Figure 3B**).

**Figure 3.**
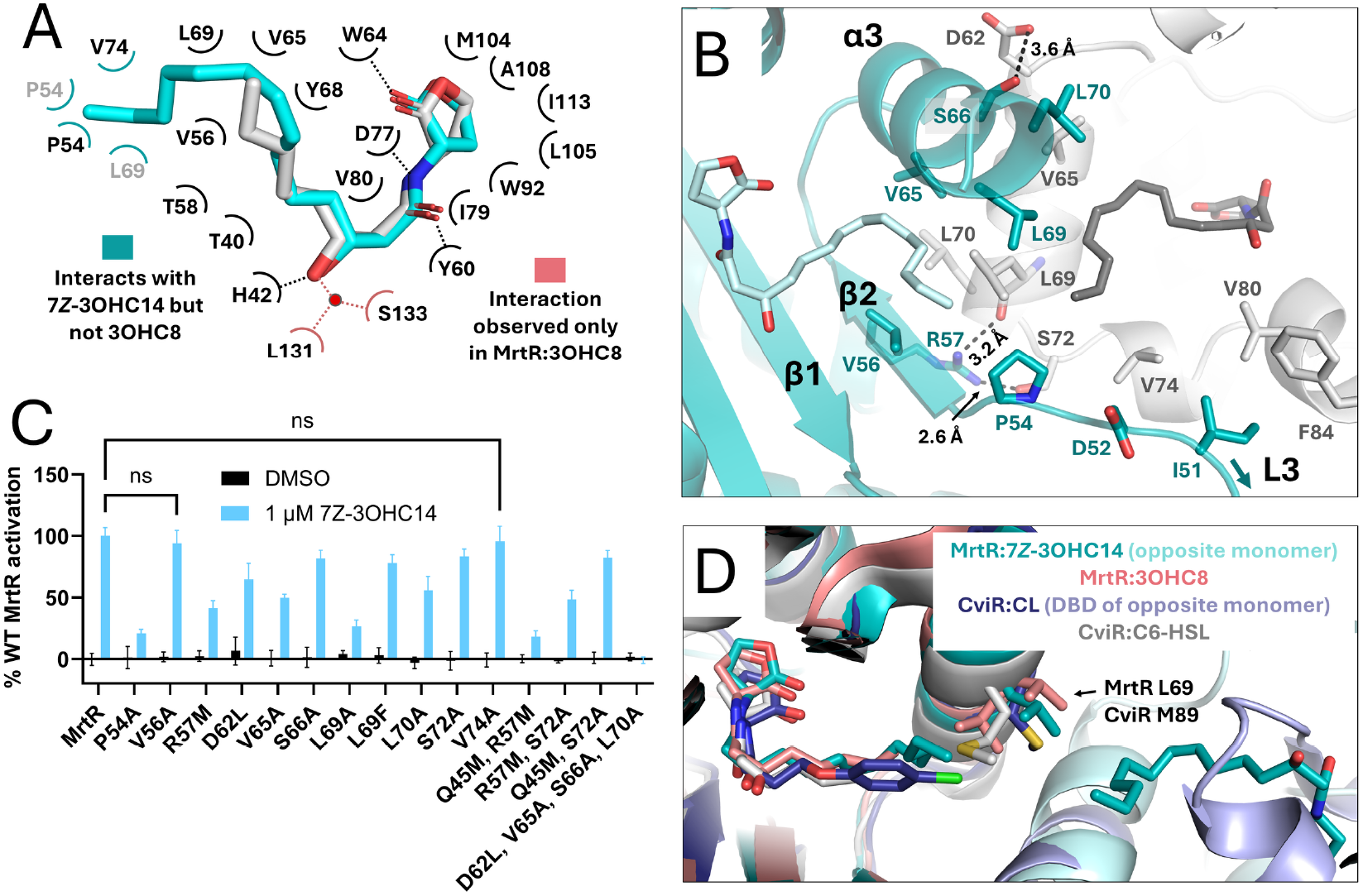
Contacts affecting MrtR agonism and ligand specificity. **(A)** Schematic showing residues that interact with ligand in MrtR:3OHC14 and MrtR:3OHC8. Gray labels indicate residues from the opposite monomer. The water molecule that mediates hydrogen bonding from the AHL 3OH to L131 and S133 is not resolved in MrtR:7*Z*-3OHC14, possibly due to the resolution of this structure. **(B)** Interactions at the dimerization interface near the ligand binding pocket in MrtR:7*Z*-3OHC14. Potential hydrogen bonds shown in black with distances indicated. All side chains shown can engage either in intermonomer hydrogen bonds or hydrophobic interactions (3.6-4.3 Å) with the ligand or with the opposite monomer. See **Figure 2C** for contacts to D52. **(C)** Transcriptional activation of point mutants of residues shown in panel **(B)** in an *E. coli* MrtR reporter. Unless indicated as “ns,” mutants have maximum activity significantly lower than WT MrtR based on an ANOVA with Dunnett’s correction for multiple comparisons and p ≤ 0.01. Data represent at least three biological replicates, each performed in technical triplicate. Error bars represent standard deviation. **(D)** Comparison of CviR and MrtR bound to agonist vs. antagonist. Note that the MrtR dimerization interface occupies the same face of the LBD as the CviR DBD in its CL-bound “crossed-domain” conformation. The conformation of MrtR L69 changes in structures bound to agonist (7*Z*-3OHC14) vs. antagonist (3OHC8). The analogous residue in CviR, M89, likewise changes position in structures bound to agonist (C6-HSL; PDB ID: 3QP1) vs. antagonist (CL; PDB ID: 3QP5) (13).

The activity of MrtR mutants in *E. coli* reporters demonstrated the importance of dimerization contacts along helix α3, strand β2, and loop L3 for MrtR activation. L69A and P54A caused the largest decreases in activity (∼75% and ∼80%, respectively), while L69F lost only ∼20% activity, consistent with the central role of these residues in mediating MrtR activation. Mutations to R57, D62, V65, S66, L70, or S72 showed intermediate (∼20-60%) decreases in activity individually, and simultaneous mutation of multiple dimerization contacts (R57M/Q45M or D62L/V65A/S66A/L70A) showed an additive effect (**Figure 3C**).

Residue L69 changes conformation based on bound ligand, pointing out of the ligand-binding pocket to interact with the opposite monomer in agonist-bound MrtR and pointing in towards the ligand and displaying conformational heterogeneity in antagonist-bound MrtR (**Figures S13** and **S14**). Interestingly, MrtR L69 corresponds to CviR M89, which also undergoes a conformational shift in response to agonist (C6-HSL) vs. antagonist (C10-HSL or CL) binding and is believed to play a crucial role in stabilizing the LBD-DBD interaction in antagonist-bound CviR (13). The dimerization interface in agonist-bound MrtR occupies the same face of the LBD as the LBD-DBD interface in antagonist-bound CviR (**Figure 3D**), and when L69 or M89 points out of the binding site, it promotes interdomain interaction (in agonist-bound MrtR or antagonist-bound CviR). As a result, MrtR L69 and CviR M89 play analogous roles in these receptors by shifting in towards the ligand or out towards an interacting domain in response to the AHL acyl tail length and stabilizing interdomain interactions that confer either an active or inactive receptor conformation.

### Proposed mechanism of MrtR activation and inhibition

The biochemical and structural data above lead us to a molecular model for small molecule-mediated agonism and antagonism of MrtR. The 14-carbon ligand tail of agonists can interact with hydrophobic residues along helix α3, strand β2, and adjacent to loop L3, stabilizing intermonomer contacts and promoting dimerization. The conformation of L3 may additionally affect the DBD conformation through the hydrophobic pocket formed by I47 of the LBD, V176 of the linker, and L236 of the DBD, thereby promoting a DBD conformation that is poised for DNA binding. In contrast, antagonists can bind to MrtR and displace its native agonist, but the shorter 8-carbon tail is less able to engage in key contacts near the dimerization interface and L3.

Compound screening data (**Figure 1B**) support that tail length plays an important role in MrtR agonism vs. antagonism: as the tail length of native AHLs increases from 8 to 14 carbons, antagonism decreases and agonism increases. The most active synthetic antagonist (H23) has an acyl tail of 8 atoms in length, analogous to highly active native AHL antagonists. In addition, structural data support the enhanced ability of 14-carbon agonists vs. 8-carbon antagonists to stabilize important dimerization elements (loop L3 and residue L69), as these elements are relatively well-defined in the lower-resolution (> 3 Å) agonist structures but have poor density in the higher-resolution (< 1.5 Å) antagonist structures. L3 and residue L69 are better defined in MrtR bound to 3OC8+TEG than in the 3OC8 and 3OHC8 antagonist structures (**Figures S8** and **S14**), possibly because the TEG contacts residues that interact with the 14-carbon agonists but not the 8-carbon antagonists (e.g., P54).

## DISCUSSION

The structures presented here provide, to our knowledge, the first full-length views of a LuxR-type receptor bound to both an agonist and an antagonist and establish a framework for understanding MrtR activation and inhibition. In contrast to many well-characterized LuxR-type receptors that are regulated by ligand-induced changes in solubility or folding (14, 27, 39-41), MrtR activity appears to be primarily determined by its dimerization state, with long AHL tails stabilizing intermonomer interactions.

Comparison with LasR and QscR reveals a shared mechanistic feature among receptors that preferentially bind long-chain AHLs (≥ 12 carbons) — a central role for loop L3 in ligand response and receptor activation. In these systems, the ligand extends toward residues near both the dimerization interface and L3. Consistent with this positioning, L3 contributes to dimerization (LasR (14), QscR (36), and MrtR) and/or interdomain contacts (QscR and MrtR), and its conformation varies as a function of the bound ligand (LasR (37, 42) and MrtR). These observations support a model in which ligand-dependent modulation of L3 contributes to coupling ligand binding to oligomerization and activation. In contrast, receptors that preferentially bind shorter AHLs (≤ 8 carbons), including SdiA (31), RhlR (32), and TraR (34), also exhibit L3-mediated hydrophobic interdomain contacts but position the ligand farther from L3. This spatial separation likely reduces the extent to which ligand binding directly influences L3 conformation, suggesting a mechanistic distinction between long- and short-chain AHL sensors. Additional structural and biochemical studies across LuxR-type receptors with diverse ligand preferences will be important to determine the generality of this model.

Our findings also highlight both the value and current limitations of protein structure prediction for understanding LuxR-type receptor function. The AlphaFold (29) model of apo-MrtR closely resembled the experimentally determined agonist-bound structure (**Figure S4**) and proved useful here for molecular replacement, but it did not provide insight into ligand-dependent conformational changes. Likewise, ligand-inclusive predictions generated using the Chai Discovery platform (43) yield nearly identical conformations for MrtR in agonist- and antagonist-bound states (**Figure S15**), in contrast to the distinct structural differences observed experimentally. These discrepancies underscore the importance of experimental structural and biochemical approaches for defining ligand-response mechanisms in this receptor family.

Finally, these results may have broader relevance for QS in rhizobia. MrtR homologs are highly conserved in several agriculturally important species (**Table S1**), including CinR in *Rhizobium etli* and *R. leguminosarum*, which share > 95% sequence identity with MrtR and have established roles in nodule formation, nitrogen fixation (22), and symbiotic plasmid transfer (19). In addition, rhizobia and other soil-dwelling bacteria (24, 44) produce AHLs that we show here to strongly modulate MrtR activity, which could contribute to interspecies interactions. Thus, the structural insights reported here may extend beyond MrtR, contributing to our understanding of LuxR-type receptor function in ecologically important systems.

## METHODS

Standard reagent and instrumentation information and additional details on the methods below are provided in SI appendix.

### Compound handling

Compounds used for screening were purchased from Sigma-Aldrich, Chemodex, or Cayman Chemical or acquired from in-house stocks synthesized according to published methods (**Table S3**). Compounds 3OHC8, 3OHC14, and 7*Z*-3OHC14 were a mix of 3*R* and 3*S* diastereomers, while 3*R*-OHC12 and 3*S*-OHC12 were enantiopure (26).

### Compound and mutant activity testing

Reporter assays were performed as previously described (25). See **Table S8** for strains and plasmids and **Table S9** for primers.

**Fluorescence polarization, chemical crosslinking, and differential scanning fluorimetry** were performed according to standard methods.

### Crystal structure determination and analysis

MrtR was crystalized using sitting drop vapor diffusion. Structures were refined using the CCP4 suite (45). PDB ID codes for MrtR structures bound to indicated compounds are as follows: 7*Z*-3OHC14 (9Y2H), 3OHC14 (9ZZ0), 3OHC8 (9ZPJ), 3OC8 (11BM), and 3OC8+TEG (10ZI).

## Supporting information

Structure validation reports

## ACKNOWLEDGEMENTS

Financial support for this work was provided by the NIH (R35 GM131817). I.M.S. was supported in part by an NSF Graduate Research Fellowship (DGE-2137424) and is an affiliate of the UW−Madison NIH Chemistry−Biology Interface Training (CBIT) Program (T32 GM008505 and T32 GM152341). We thank Isabel Cannell for the synthesis of 3OC16, 3OC18, and 3OC20; the laboratories of Lloyd Smith and Samuel Gellman for instrumentation; Craig Bingman at the UW– Madison Crystallography Core; and Mikael Elias and Andrew Buller for technical advice.

