## Supporting Information and Data for "MrtR of *Mesorhizobium tianshanense* reveals both activation and inhibition mechanisms of a LuxR-type quorum sensing receptor"

### CONTENTS

- Supplemental methods
- **Figure S1.** Activity of synthetic AHL analogs in an *E. coli* MrtR reporter.
- **Figure S2.** Dose-response curves for compounds in an *E. coli* MrtR reporter, corresponding to data in **Table 1**.
- **Figure S3.** SDS-PAGE data for chemical crosslinking of MrtR.
- **Figure S4.** Comparison of MrtR:7Z-3OHC14 crystal structure and AlphaFold model.
- **Figure S5.** Comparison of MrtR bound to 7Z-3OHC14 vs. 3OHC14.
- **Figure S6.** Alignment of MrtR domains with other structures of LuxR-type receptors.
- **Figure S7.** Dimerization interface comparison of MrtR:7Z-3OHC14 and other structures of LuxR-type receptors.
- **Figure S8.** L3 loop densities for MrtR structures.
- **Figure S9.** Ligand binding poses in MrtR and other receptors.
- **Figure S10.** Residue H42 affects MrtR's preference for AHL oxidation state.
- **Figure S11.** Ligand binding poses in TraR:3OC8, MrtR:3OC8, and MrtR:3OHC8.
- **Figure S12.** Antagonism and agonism activity of 3OC8 in MrtR mutants.
- **Figure S13.** Conformational change for L69 between MrtR agonist- and antagonist-bound structures.
- **Figure S14.** Density for L69 in MrtR structures.
- **Figure S15.** Chai Discovery predictions of MrtR structure in the presence of 7Z-3OHC14 or 3OC8.
- **Figure S16.** Representative FPLC trace from MrtR purification.
- **Figure S17.** Representative SDS-PAGE gel from MrtR purification.
- **Figure S18.** Omit maps showing electron density calculated without the AHL modeled.

- **Table S1.** Abbreviated table of BLAST results for MrtR.
- **Table S2.** Abbreviated table of BLAST results for the promoter sequence in the *E. coli* MrtR transcriptional reporter.
- **Table S3.** Origins of compounds tested in this study.
- **Table S4.** Molecular weight of MrtR complexes estimated based on size exclusion chromatography.
- **Table S5.** X-ray crystallographic data collection and refinement statistics
- **Table S6.** Assemblies predicted for MrtR:7Z-3OHC14 by PDBePISA.
- **Table S7.** Assemblies predicted for MrtR:3OHC14 by PDBePISA.
- **Table S8.** Bacterial strains and plasmids used in this study.
- **Table S9.** Primers and oligos used in this study.
- Relative fluorescence unit (RFU) and first derivative traces for MrtR DSF.
- Size exclusion chromatography traces used to estimate MrtR complex molecular weight.
- References

<u>**Supplemental methods.**</u>

**General reagent and instrumentation information.** All standard reagents and solvents were purchased from commercial sources and used without purification. Arabinose was purchased from MilliporeSigma; Tris(2-carboxyethyl)phosphine hydrochloride (TCEP-HCl) from Sigma-Aldrich; ampicillin, kanamycin, gentamycin, and isopropyl $\beta$-D-1-thiogalactopyranoside (IPTG) from Dot Scientific; and dithiothreitol (DTT) from Tokyo Chemical Industry (TCI). Water (18 MΩ) was purified with a Sartorius Arium Pro system. Antibiotic, DTT, and TCEP stocks were prepared at 1000x in water, stored at -20 °C, and added to buffer or media immediately before use. Glucose was prepared as a 50% solution in water, filtered with Steriflip vacuum filter units (Millipore), and stored at 4 °C. DNA oligos used for cloning and in DNA-binding assays were purchased from Integrated DNA Technologies (IDT) and stored at -20 °C after resuspension in water. Fluorescence and $OD_{600}$ data were collected on a BioTek Synergy 2 plate reader using Gen5 software (version 3.12). Arabinose stocks were prepared in water and stored at room temperature for up to 6 months.

**Compound handling.** Compounds used for screening were purchased from Sigma-Aldrich, Chemodex, or Cayman Chemical or acquired from in-house stocks synthesized according to published methods (see **Table S3** for compound sources). Compounds 3OC16, 3OC18, and 3OC20 were synthesized through DCC/DMAP-catalyzed coupling of an appropriate aliphatic carboxylic acid to Meldrum's acid, followed by a subsequent amide coupling to the L-homoserine lactone head group, as previously reported (1). Compound stock solutions were prepared in DMSO and stored at -20 °C.

**Bacteriology.** The bacteria and plasmids used in this study are listed in **Table S8,** and the primers are listed in **Table S9**. Cultures were grown in Lennox Broth (LB) (Research Products International) with appropriate antibiotics and incubated at 37 °C with shaking (200 rpm). Strains were stored in 25% glycerol at -80 °C and streaked on LB agar plates with appropriate antibiotics. Chemically competent cells used for cloning were prepared in-house.

**Plasmid and reporter construction.** MrtR mutants were created using the Q5 Site-Directed Mutagenesis Kit according to the manufacturer's protocol (New England Biolabs cat. no. E0554S) using pJN105-mrtR as a template. Mutations were verified by sequencing, and the resulting plasmids were transformed into *E. coli* BW27749 with pmrtI:GFP to create transcriptional reporters.

For pET28-his6-SUMO-mrtR, the vector was amplified from pET28-his6-SUMO-smaR, the insert was amplified from pJN105-mrtR, and the fragments were assembled by Gibson assembly (New England Biolabs cat. no. E2611). Insertion was verified by sequencing, and the resulting plasmid was transformed into *E. coli* BL21(DE3).

**Transcriptional reporter assays.** Reporter assays were performed as previously described with minor modifications (2). Overnight cultures of reporter strains were diluted 1:10 in fresh LB medium with 100 µg/mL ampicillin and 15 µg/mL gentamicin. Cultures were grown to an $OD_{600}$ of 0.22-0.27 as measured in a 96-well microtiter plate with 200 µL culture per well. Aliquots (2 µL) of compound stocks or DMSO (vehicle) were added to each well of black 96-well plates with transparent bottoms (Corning 3904). Cultures were induced by adding 1:100 of a 25 µg/mL stock of arabinose (for a 0.25 µg/mL final concentration), and 198 µL of culture was added to each well. Uninduced culture served as a control. Data normalized to WT MrtR saturated with 7Z-3OHC14 used 1-10 µM 7Z-3OHC14 (see **Figure S10** for dose-response curve). Plates were incubated at 37 °C with shaking at 200 rpm for 3.5 h. Plates were mixed by pipetting before reading GFP fluorescence and $OD_{600}$. Microsoft Excel (version 16.92) was used to normalize data, and GraphPad Prism (version 10.6.1) was used to generate graphs and perform statistical analyses. ANOVA was used to compare activity across mutants with a Dunnett correction for multiple comparisons.

**MrtR purification.** MrtR was expressed in *E. coli* BL21(DE3) with an N-terminal His6-SUMO tag. The strain was maintained with 50 µg/mL kanamycin throughout growth and expression. A fresh colony was used to inoculate 1 L of LB medium supplemented with 0.75% glucose in a 4 L flask. Culture was grown at 37 °C with shaking (200 rpm) to an $OD_{600}$ of 0.6-0.8 and then chilled on ice for 10 min. Protein production was induced with 0.2 mM IPTG before incubation for ~16 h at 17 °C with shaking (200 rpm). Cells were chilled on ice, harvested by centrifugation, resuspended in 50 mL of buffer (25 mM HEPES, 200 mM NaCl, 50 mM imidazole, 1 mM DTT, pH 7.4), and lysed by sonication. Clarified lysate was filtered and loaded onto a 5 mL HisTrap HP column (Cytiva) using an ÄKTA chromatography system (Cytiva). MrtR was eluted with a gradient of 50-360 mM imidazole (**Figure S16**), then dialyzed against buffer without imidazole (25 mM HEPES, 150 mM NaCl, 1 mM DTT, pH 7.4) for 4 h at 4 °C. The His6-SUMO tag was cleaved with Ulp1 (TriAltus Bioscience) overnight at 4 °C while the protein continued to dialyze against fresh buffer. Cleaved tag and protease were removed with HisPur Ni-NTA or cobalt resin (Thermo Scientific), and purified MrtR was concentrated to 120-250 µM using Amicon Pro 50 kDa cutoff centrifugal filters (Millipore Sigma). Protein concentration was estimated based on A280 measured on a Nanodrop 2000c spectrophotometer (Thermo Scientific). Aliquots of MrtR in 0.2 mL PCR tubes (Dot Scientific) were frozen in liquid nitrogen and stored at -80 °C and thawed on ice before use. See **Figure S17** for SDS-PAGE gel from purification.

**SDS-PAGE.** Samples were prepared with 12 µL protein sample, 4.5 µL of 4x LDS buffer (G-Biosciences cat. #786323), and 1.5 µL of 20 mM DTT, then heated to 95 °C for 10 min before loading on a 10% Mini-PROTEAN TGX gel (Bio-Rad) or a hand-cast 12.5% polyacrylamide gel. Gels were run on ice for 100 min at 120V. Gels were stained with Coomassie Brilliant blue R-250 (Santa Cruz Biotechnology) and imaged using an iPhone X or a Cytiva Amersham Typhoon imager.

**Fluorescence polarization (FP).** A FAM-labeled 42-bp section of the *mrtI* promoter (3) was used for fluorescence polarization measurements. The same sequence without a FAM label was used as a specific competitor, and a 32-bp section of pUC18 was used as a nonspecific competitor (4) (see **Table S9** for sequences). DNA was diluted to 1 µM and annealed by heating to 95 °C and cooling slowly. MrtR was diluted to 1600 nM in binding buffer (20 mM HEPES, 5 mM $MgCl_2$, 80 mM NaCl, 5% glycerol, pH 7.0, 5 mM DTT) with ligand of interest or 1% DMSO. Samples were prepared in a 384-well black microtiter plate (Corning 3575) with final concentrations of 400 nM MrtR, 100 µM ligand, and 5 nM FAM-labeled DNA probe. Control samples contained 500 nM nonspecific or specific competitor DNA. Samples were incubated on ice for 15 min before reading fluorescence polarization on a PerkinElmer EnVision 2105 multimode plate reader. Microsoft Excel (version 16.92) was used to normalize data, and GraphPad Prism (version 10.6.1) was used to generate graphs and perform statistical analyses. An ANOVA was used to compare activity between conditions with a Dunnett correction for multiple comparisons. Outliers were identified and removed using a ROUT test with Q = 1%.

**Chemical crosslinking.** MrtR was diluted to 5 µM in buffer (25 mM HEPES, 150 mM NaCl, 2 mM DTT, pH 7.4) and incubated with 100 µM compound or 1% DMSO on ice for 10-30 min before the addition of 0.375 mM $BS(PEG)_5$ (Sigma-Aldrich). Samples were statically at room temperature for 1 h, then quenched with 10% v/v of 1 M Tris buffer (pH 7.5) for 15 min before preparing samples for SDS-PAGE.

**Differential scanning fluorimetry (DSF).** MrtR was diluted to 5 µM in buffer (25 mM HEPES, 150 mM NaCl, 1 mM DTT, pH 7.4), and incubated with 100 µM ligand or DMSO on ice for 25 min. Samples were centrifuged at 15,000 x g for 3 min, mixed 1:1 with 10x SYPRO orange protein stain (Sigma-Aldrich) in a white PCR plate (Bio-Rad MLP9651), and heated from 20–95 °C at 1 °C/min on a C1000 Touch Thermal Cycler with a CFX96 Real-Time detection system to measure fluorescence. DSFworld (5) was used to calculate $T_m$ values for all conditions except for DMSO, for which the melting curves were too shallow for consistent automatic determination of $T_m$, so $T_m$ values were manually determined based on the minimum -dRFU/dt value. Raw fluorescence and first derivative curves are provided at the end of this SI Appendix.

**Size exclusion chromatography (SEC).** MrtR was diluted to 20 µM in buffer (25 mM HEPES, 150 mM NaCl, 5 mM TCEP, pH 7.4) with ligand of interest or DMSO (100 µM 3OC8, 3OC12, or 7*Z*-3OHC14, 50 µM C14-HSL, all at 1% DMSO), and ~75 µL was loaded onto a Superdex 200 Increase 10/300 column. Separation was performed at 4 °C at a flow rate of 0.3 mL/min with a mobile phase of 25 mM HEPES, 150 mM NaCl, 5 mM TCEP, pH 7.4 supplemented with the ligand of interest or DMSO (100 µM of 3OC8, 10 µM 3OC12, 5 µM C14-HSL, or 1 µM 7*Z*-3OHC14, all at 0.1% DMSO). Fresh TCEP (pH ~7.5) was added to buffer daily. A standard curve was constructed using bovine serum albumin (BSA) (Research Products International A30075), ovalbumin (Sigma-Aldrich A5503), carbonic anhydrase (Sigma-Aldrich C7025), and equine cytochrome C (Sigma-Aldrich C2506) and used to estimate the molecular weight of MrtR peaks. Standards were run in the presence of DMSO, 3OC12, and 7*Z*-3OHC14 and showed nearly identical retention times in each condition, so standard runs were pooled to create the standard curve. Full traces are shown at the end of this SI Appendix, including a mock injection where a sample was prepared in the same manner as the protein samples (with DMSO added) but with no protein and a blank where only running buffer was injected. Peak retention time was determined using UNICORN control software (version 5.2, General Electric Company).

**Crystallization of MrtR.** MrtR bound to 7*Z*-3OHC14 was crystallized in 50 mM sodium HEPES, 50 mM MOPS, pH 7.5; 30 mM $MgCl_2$, 30 mM $CaCl_2$, 30% ethylene glycol, 10% PEG 8000, and ~1% DMSO. MrtR bound to 3OHC14 was crystallized in 50 mM imidazole, 50 mM MES, pH 6.5; 30 mM $MgCl_2$, 30 mM $CaCl_2$, 30% ethylene glycol, 5% PEG 8000, and ~1% DMSO. MrtR bound to 3OC8 or 3OHC8 was crystallized in 0.05 M imidazole, 0.05 M MES, pH 6.5; 0.03 M each of diethylene glycol, triethylene glycol, tetraethylene glycol, and pentaethylene glycol; 12.5% v/v racemic 2-methyl-2,4-pentanediol (MPD); 12.5% PEG 1000; 12.5% w/v PEG 3350 (condition 2-4 from Morpheus screen MD1-46 from Molecular Dimensions) with the addition of 300–1500 µM ligand of interest and ~3% DMSO. Protein at 120–250 µM was mixed with crystallization buffer in sitting drop plates (Art Robbins 102-0003-00 or Hampton HR1-002). Plates were incubated at 4 °C, and crystals appeared in 1–3 days. Crystals grew in absence of ligand in the same conditions used for 3OC8 and 3OHC8, as well as in 50 mM imidazole, 50 mM MES, pH 6.5; 0.12 M each of diethylene glycol, triethylene glycol, tetraethylene glycol, and pentaethylene glycol; 12.5% v/v MPD; 12.5% PEG 1000; 12.5% w/v PEG 3350, but the isolated apo-MrtR crystals produced poor-quality data with evidence of twinning. Crystals with 7*Z*-3OHC14 or 3OHC14 grew best using a 90–120 µM MrtR stock, while crystals with 3OC8 or 3OHC8 grew best with a ~245 µM MrtR stock.

Diffraction data were collected at the National Synchrotron Light Source II (NSLS-II) beamline 17-ID-2 for the 7*Z*-3OHC14 and 3OHC14 datasets and at the European Synchrotron Radiation Facility (ESRF) beamline BM07 for 3OHC8 and beamline MASSIF-I for 3OC8 datasets. Structures were solved by molecular replacement using Phaser (6) based on the AlphaFold model for MrtR (**Figure S4**), and the model was built using Coot (7) and refined using Refmac5 (8) in the CCP4 program suite (9). FreeRflag (10) and Matthews_coef (11) programs were also used. The 3*R* diastereomers of 7*Z*-3OHC14 and 3OHC8 provided the best fit to the density in these structures (mixtures of 3*S* and 3*R* diastereomers used in crystallizations). The density in the 3OHC14 structure did not clearly indicate one diastereomer over the other, so the 3*R* diastereomer was modeled. The PDB extract (12) web server was used to prepare coordinate files for deposition (13) to the Worldwide Protein Data Bank (wwPDB.org) (14). Protein interfaces, surfaces, and assemblies service (PISA) v 1.52 at the European Bioinformatics Institute (15) was used to predict assembly state. TLS refinement was used for the 7*Z*-3OHC14 and 3OHC14 datasets, and anisotropic refinement was used for 3OHC8 and both 3OC8 datasets. See **Table S5** for data collection and refinement statistics and **Figure S18** for omit maps for the ligand-binding site. Figures of the structures were created using PyMOL (The PyMOL Molecular Graphics System, version 3.1.4.1, Schrödinger, LLC.).

**Structure predictions.** The AlphaFold 3 (16) web server was used to predict the structure of MrtR (**Figure S4**). The Chai Discovery (17) web server was used to predict MrtR structures with 7*Z*-3OHC14 or 3OC8, using the Multiple Sequence Alignments and Templates options (**Figure S15**).

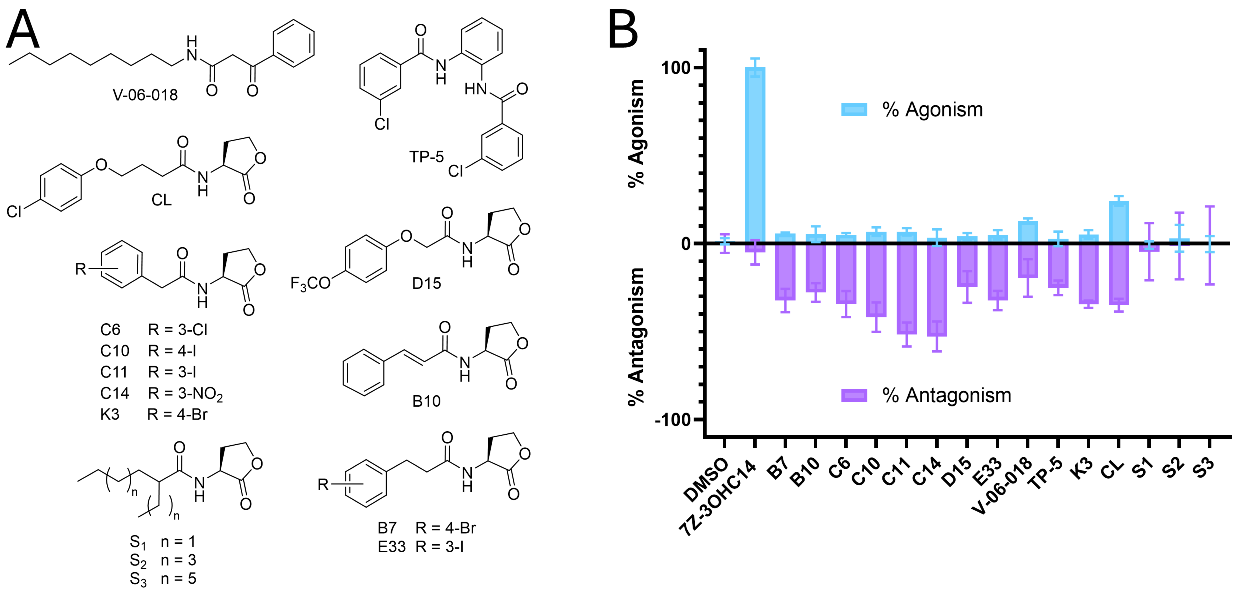


**Figure S1. Activity of synthetic AHL analogs in an *E. coli* MrtR reporter. (A)** Structures of synthetic compounds assayed for activity at MrtR. **(B)** Agonism and competitive antagonism activity of compounds in an *E. coli* MrtR transcriptional reporter. Activity of DMSO and 7*Z*-3OHC14 is shown for reference. Percent agonism represents transcription of *mrtI*:GFP with each compound at 10 µM, normalized to uninduced culture (0%) and MrtR saturated with 7*Z*-3OHC14 (100%). Percent antagonism represents transcription of *mrtI*:GFP with 10 µM of each compound competed against 8 nM 7*Z*-3OHC14, normalized to uninduced culture (-100%) and 8 nM 7*Z*-3OHC14 (0%). Data represent at least one biological replicate performed in technical triplicate; activity for compounds with > 10% agonism or > 40% inhibitory activity was confirmed with an additional biological replicate or a dose-response curve. Error bars represent standard deviation.

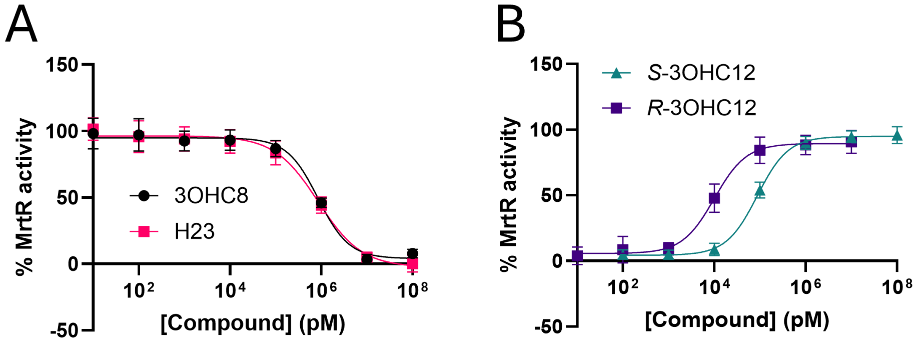


**Figure S2. Dose-response curves for compounds in an *E. coli* MrtR reporter, corresponding to data in Table 1. (A)** Antagonism. Data are normalized to uninduced culture (0%) and 8 nM 7*Z*-3OHC14 (100%) and represent two biological replicates, each performed in triplicate. **(B)** Agonism. Data are normalized to uninduced culture (0%) and 1 µM 7*Z*-3OHC14 (100%) and represent at least three biological replicates, each performed in triplicate.
**(All graphs)** Error bars represent standard deviation.

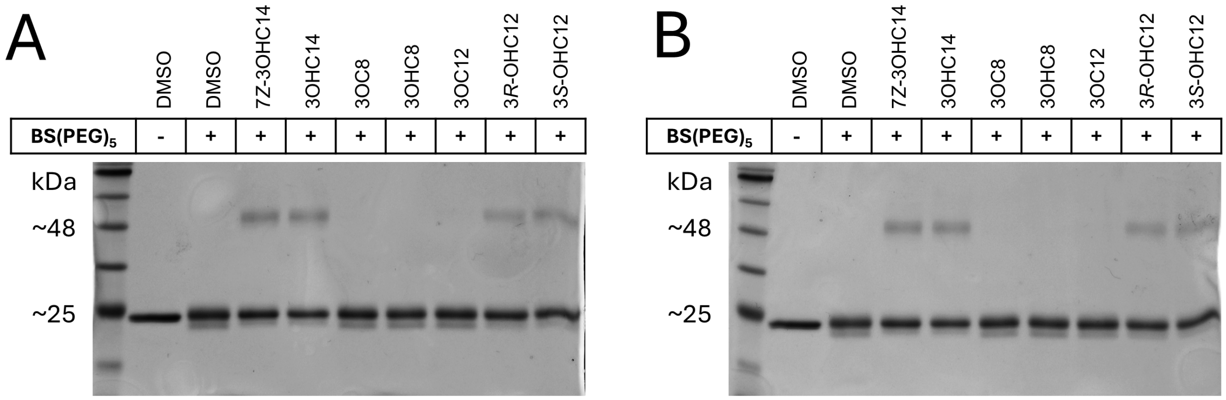


**Figure S3. SDS-PAGE data for chemical crosslinking of MrtR.** Assay performed with 5 µM purified MrtR, 100 µM compound of interest, and 0.375 mM $BS(PEG)_5$. Panels **(A)** and **(B)** represent replicate experiments. Monomeric MrtR appears just below the 25 kDa ladder band, and crosslinked bands appear just above the 48 kDa ladder band.

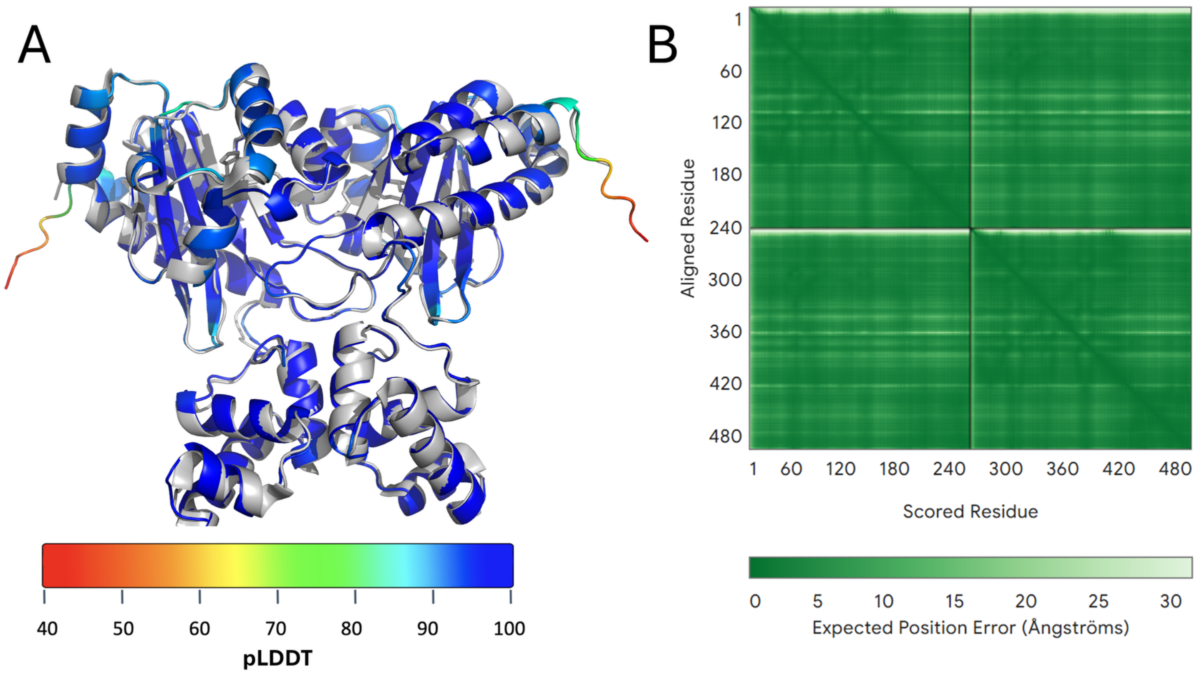


**Figure S4. Comparison of MrtR:7Z-3OHC14 crystal structure and AlphaFold model.**
**(A)** Crystal structure of MrtR:7*Z*-3OHC14 (gray) aligned with AlphaFold prediction of MrtR dimer (colored by pLDDT, a measure of prediction confidence where > 90 indicates very high confidence, 70-90 indicates high confidence, 50-70 indicates low confidence, and < 50 indicates very low confidence). Predicted and experimental structures align with RMSD of 0.54 Å.
**(B)** Plot of expected position error for AlphaFold model shown in panel **(A)**. Residues 1-241 correspond to one copy of MrtR, and residues 242-482 correspond to the second copy.

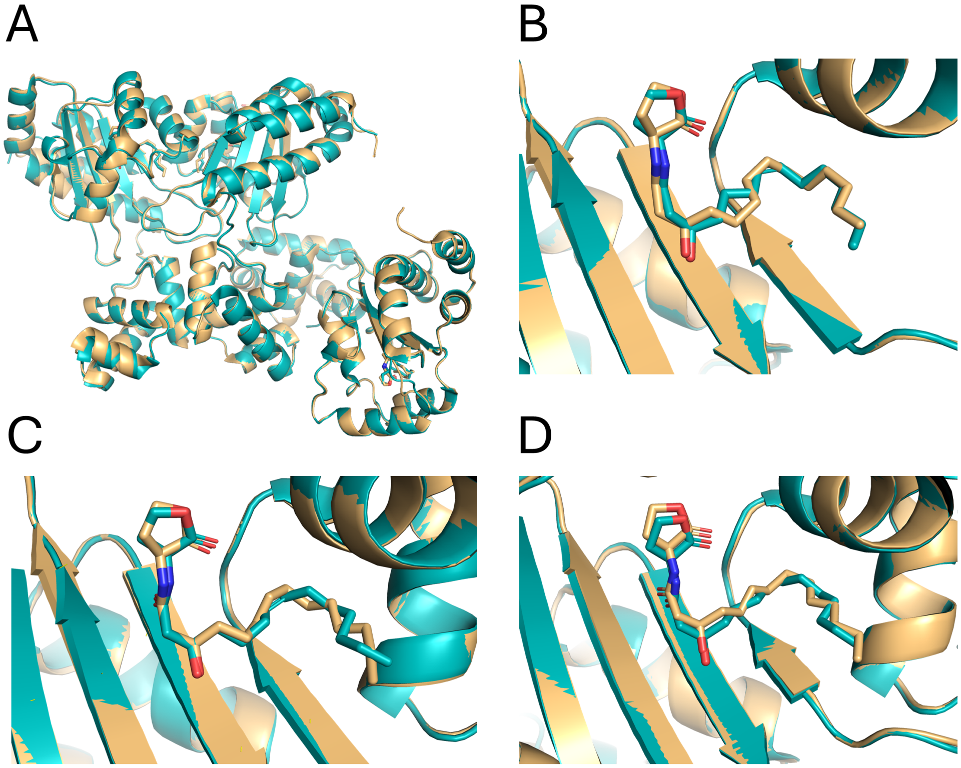


**Figure S5. Comparison of MrtR bound to 7*Z*-3OHC14 (teal) vs. 3OHC14 (tan). (A)** Image of overlayed asymmetric unit. **(B–D)** Images of overlays of the ligand-binding site of each protomer.

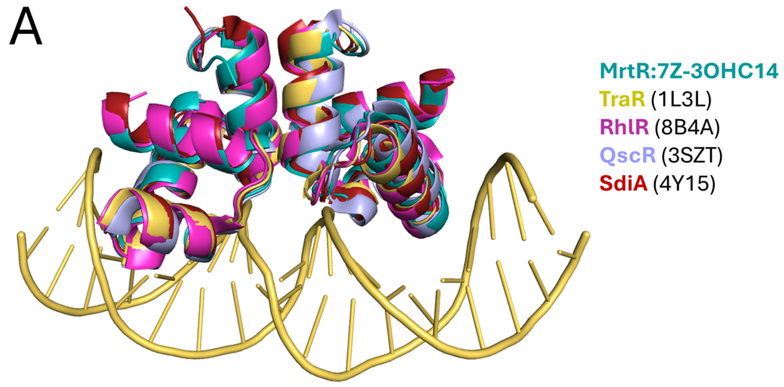


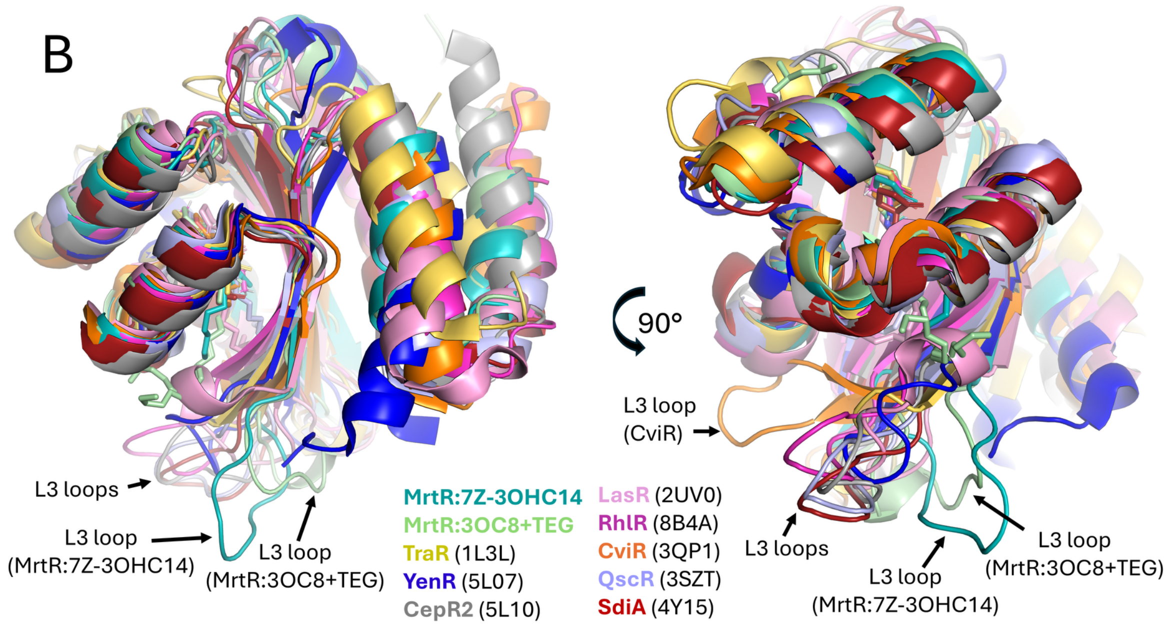


**Figure S6. Alignment of MrtR domains with other structures of LuxR-type receptors.** Alignment of **(A)** DNA-binding domains and **(B)** ligand-binding domains. Figure includes structures of TraR:3OC8 (18), YenR:apo (19), CepR2:apo (19), LasR:3OC12 (20), RhlR:C4-HSL (21), CviR:C6-HSL (22), QscR:3OC12 (23), and SdiA:3OC6 (24). DNA shown is from TraR crystal structure. PDB IDs are indicated in parentheses.

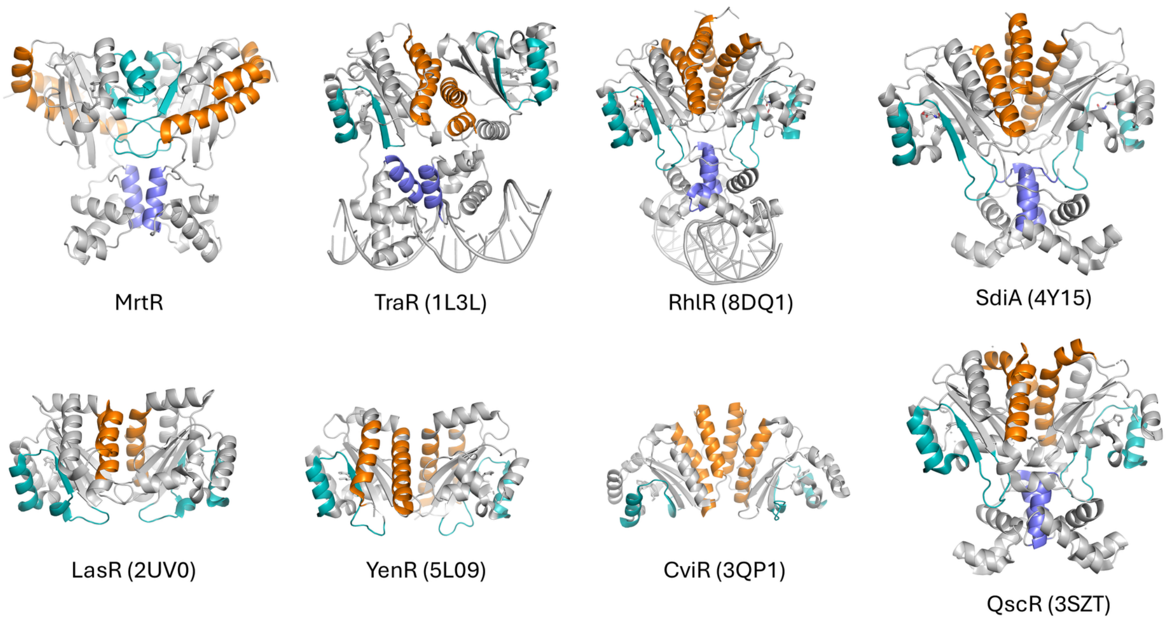


**Figure S7. Dimerization interface comparison of MrtR:7Z-3OHC14 and other structures of LuxR-type receptors.** To aid in visualization, helices α1 (res. ~5-20) and α6 (res.~145-165) of the LBD are show in orange; loop L3 (res. ~44-54), strand β2 (res. ~55-58), and helix α3 (res. ~62-71) of the LBD in teal; and the last helix of the DBD in purple. PDB IDs are indicated in parentheses for TraR:3OC8 (18), YenR:3OC6 (19), LasR:3OC12 (20), RhlR:mBTL (25), CviR:C6-HSL (22), QscR:3OC12 (23), and SdiA:3OC6 (24).

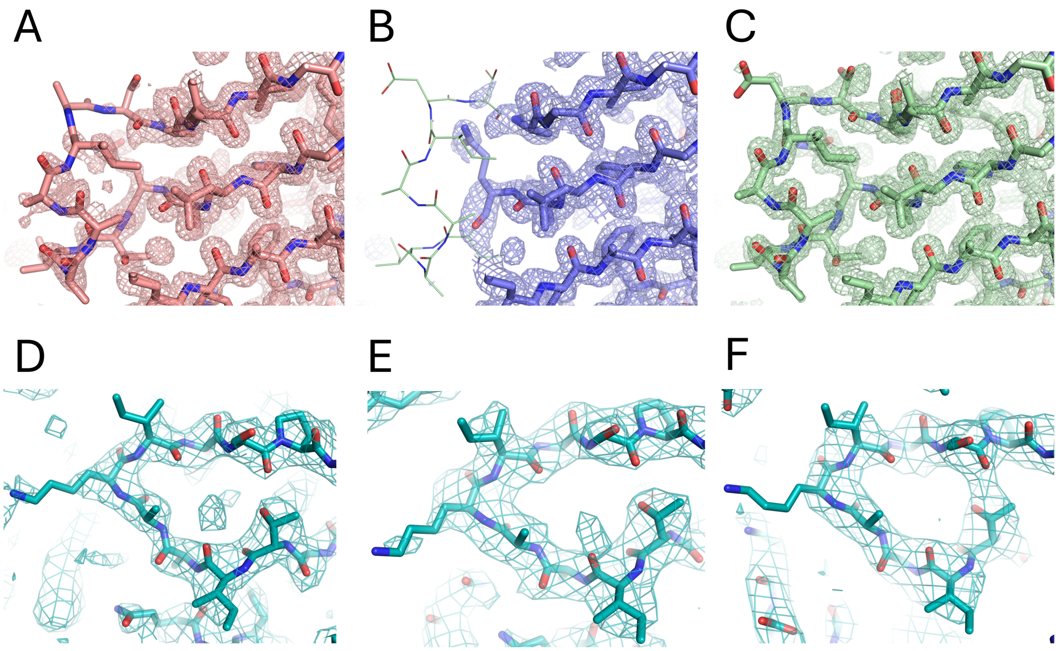


**Figure S8. Loop L3 densities for MrtR structures.** Density is shown at 1σ for MrtR bound to 3OHC8 **(A)**, 3OC8 **(B)**, 3OC8+TEG **(C)**, and all three protomers of 7*Z*-3OHC14 **(D–F)**. Green sticks in **(B)** show the position of L3 in MrtR:3OC8+TEG, as the corresponding residues are not modeled in MrtR:3OC8.

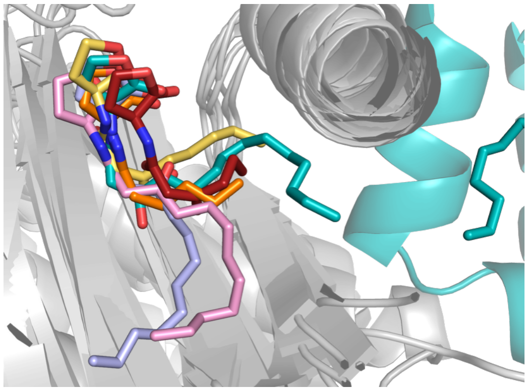

**Figure S9. Ligand binding poses in MrtR and other receptors.** Comparison of agonist-binding poses in MrtR:7*Z*-3OHC14 (teal), LasR:3OC12 (2UV0 (20), pink), CviR:C6-HSL (3QP1 (22), orange), QscR:3OC12 (3SZT (23), violet), TraR:3OC8 (1L3L (18), yellow), and SdiA:3OC6 (4Y15 (24), red). The first 160 residues of each protein (gray) were aligned, and the opposite monomer of MrtR is shown in teal.

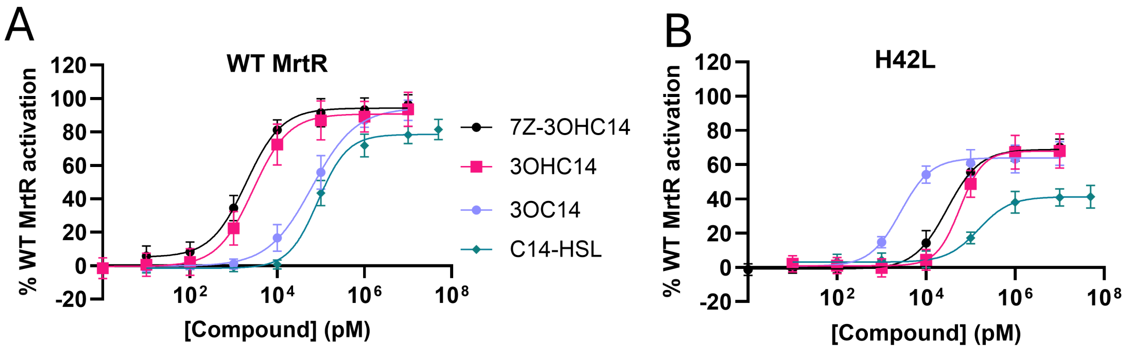


C

| **Compound** | **MrtR** | | **MrtR H42L** | |
|---|---|---|---|---|
| Agonist | $EC_{50}$ (nM) (95% CI) | Max activity (%) (95% CI) | $EC_{50}$ (nM) (95% CI) | Max activity (%) (95% CI) |
| 7Z-3OHC14 | 2.0 (1.6-2.5) | 94 (91-97) | 30 (25-36) | 69 (67-71) |
| 3OHC14 | 2.8 (2.2-3.6) | 91 (88-94) | 58 (42-78) | 68 (64-71) |
| 3OC14 | 63 (50-79) | 95 (90-99) | 2.7 (2.1-3.5) | 64 (62-66) |
| C14 | 86 (74-100) | 79 (76-81) | 150 (110-210) | 41 (39-44) |

**Figure S10. Residue H42 affects MrtR's preference for AHL oxidation state. (A)** and **(B)** Dose-response activity of AHLs with 14-carbon tails and differing oxidation states in **(A)** WT MrtR and **(B)** MrtR H42L, as measured in *E. coli* reporters. Data is normalized to uninduced culture (0%) and WT MrtR saturated with 7*Z*-3OHC14 (100%). Error bars represent standard deviation. **(C)** $EC_{50}$ values for data shown in **(A)** and **(B)**. Data for WT MrtR is also shown in **Table 1**. Data represent at least three biological replicates, each performed in technical triplicate. Maximum (Max) activity represents the top of the nonlinear fit curve based on reporter data normalized to uninduced culture (0%) and MrtR saturated with 7*Z*-3OHC14 (100%). "CI" indicates confidence interval.

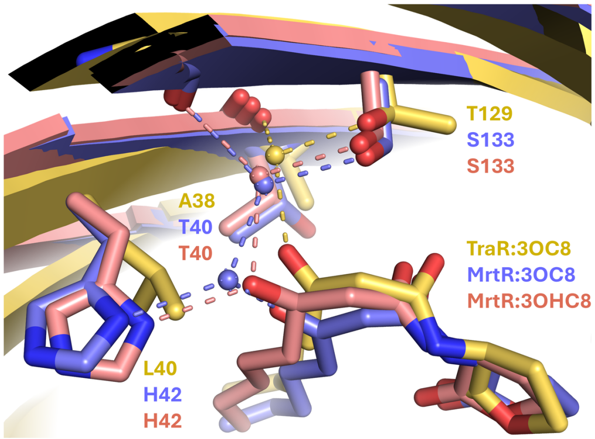


**Figure S11. Ligand binding poses in TraR:3OC8, MrtR:3OC8, and MrtR:3OHC8.** Comparison of TraR:3OC8 (1L3L (18), yellow), MrtR:3OC8 (violet), and MrtR:3OHC8 (salmon). Dashed lines represent hydrogen bonds < 3 Å. Residue H42 in MrtR corresponds to L40 in TraR, which natively responds to 3OC8. In the crystal structure of TraR, L40 appears to push the 3O group of the 3OC8 ligand into a position to hydrogen bond through a water molecule to T129 and the main chain of A38. In contrast, 3OC8 bound to MrtR is rotated towards H42. The H42L mutation in MrtR may favor a ligand binding pose more like that observed in TraR, which may explain why an H42L mutation reverses MrtR's preference for 3OH vs. 3O AHLs.

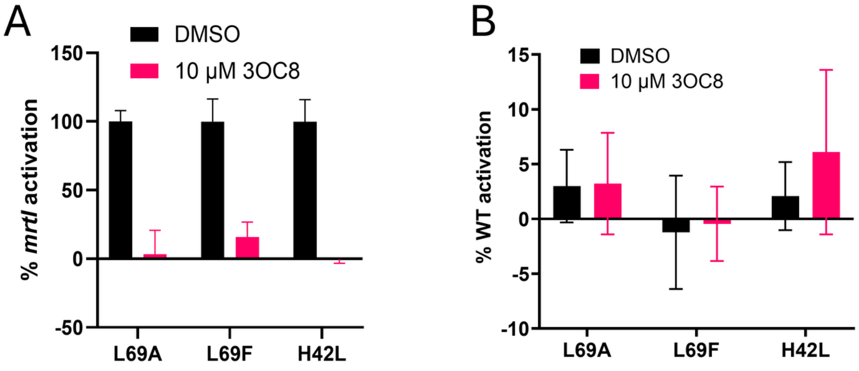


**Figure S12. Antagonism and agonism activity of 3OC8 in MrtR mutants.** Although both H42L and L69F mutations alter MrtR residues to match the identity of the corresponding residues in TraR, neither mutation converts 3OC8 from an antagonist to an agonist.
**(A)** Antagonism. Data is normalized to uninduced culture (0%) and the amount of 7*Z*-3OHC14 used for competition (100%). Approximately 4x the $EC_{50}$ value of 7*Z*-3OHC14 in each mutant was used for competition (800 nM for L69A, 24 nM for L69F, and 120 nM for H42L).
**(B)** Agonism. Data are normalized to uninduced culture (0%) and WT MrtR saturated with 7*Z*-3OHC14 (100%). **(All graphs)** Data represent two or three biological replicates, each performed in technical duplicate or triplicate. Error bars represent standard deviation.

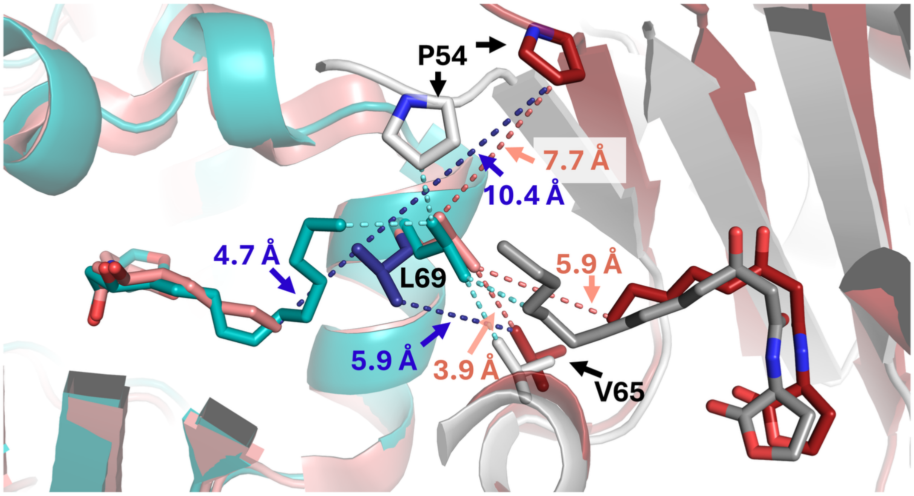


**Figure S13. Conformational change for L69 between MrtR agonist- and antagonist-bound structures.** MrtR:7*Z*-3OHC14 is shown in teal, with the opposite monomer in gray, and MrtR:3OHC8 is shown in salmon, with symmetry mate in red. In MrtR:3OHC8, L69 adopts two conformations, one pointing in towards the ligand (side chain, contacts, and distances shown in purple) and one pointing out of the ligand-binding site (side chain, contacts, and distances shown in salmon). In MrtR:7*Z*-3OHC14, L69 points out towards the dimerization interface, and distances (all 3.6–3.8 Å) are shown in teal. P54 undergoes a conformational change in agonist- vs. antagonist-bound structures and interacts with L69 in MrtR:7*Z*-3OHC14 but not in MrtR:3OHC8.

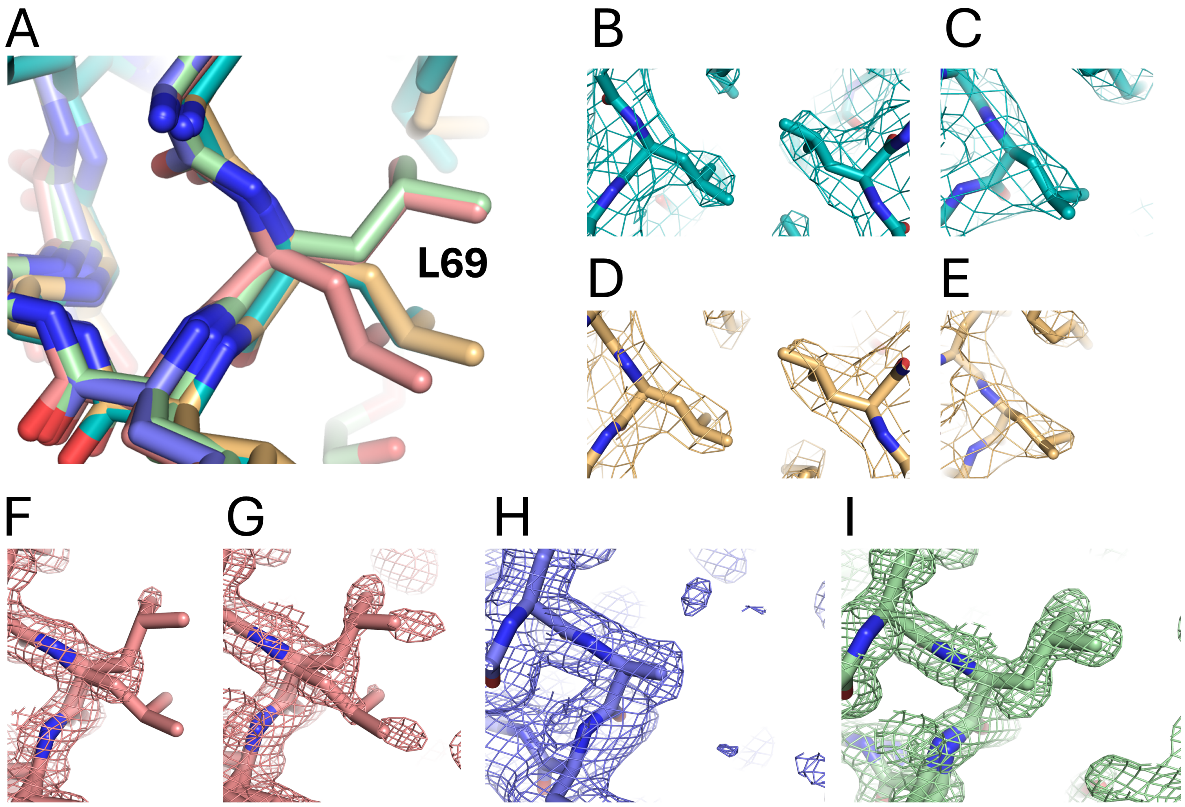


**Figure S14. Density for L69 in MrtR structures. (A)** Overlay of L69 position in MrtR:7*Z*-3OHC14 (teal), MrtR:3OHC14 (tan), MrtR:3OHC8 (salmon), MrtR:3OC8 (violet, side chain not modeled), and MrtR:3OC8+TEG (green). **(B)** and **(C)** Electron density for the three protomers of MrtR:7*Z*-3OHC14 at 1σ. **(D)** and **(E)** Electron density for the three protomers of MrtR:3OHC14 at 1σ. **(F)** and **(G)** Electron density for MrtR:3OHC8 at 1σ **(F)** and 0.5σ **(G). (H)** Electron density for MrtR:3OC8 at 0.5σ. **(I)** Electron density for MrtR:3OC8+TEG at 1σ.

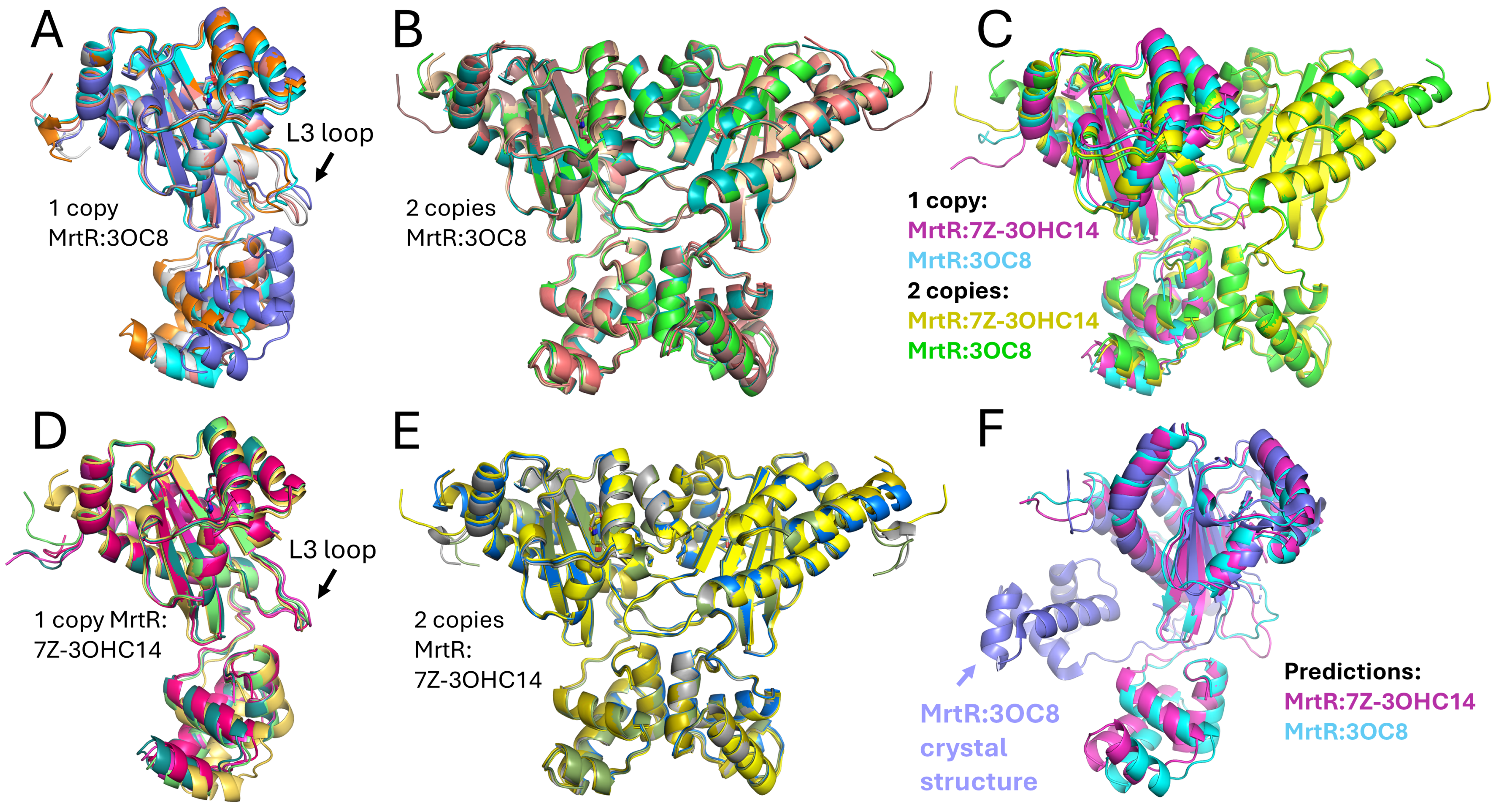


**Figure S15. Chai Discovery predictions of MrtR structure in the presence of 7Z-3OHC14 or 3OC8. (A)** Five models produced with one copy each of MrtR and 3OC8 as inputs. Note variability in loop L3 conformation. **(B)** Five models produced with two copies each of MrtR and 3OC8 as inputs. **(C)** Predicted models for two copies of MrtR:3OC8 (green), two copies of MrtR:7Z-3OHC14 (yellow), one copy of MrtR:3OC8 (blue), or one copy of MrtR:7Z-3OHC14 (magenta). **(D)** Five models produced with one copy each of MrtR and 7Z-3OHC14 as inputs. **(E)** Five models produced with two copies each of MrtR and 7Z-3OHC14 as inputs. **(F)** Predicted models for one copy of MrtR:3OC8 (blue) and MrtR:7Z-3OHC14 (magenta) aligned with crystal structure of MrtR:3OC8 (violet). Note difference in DBD orientation in predicted vs. experimental structure.

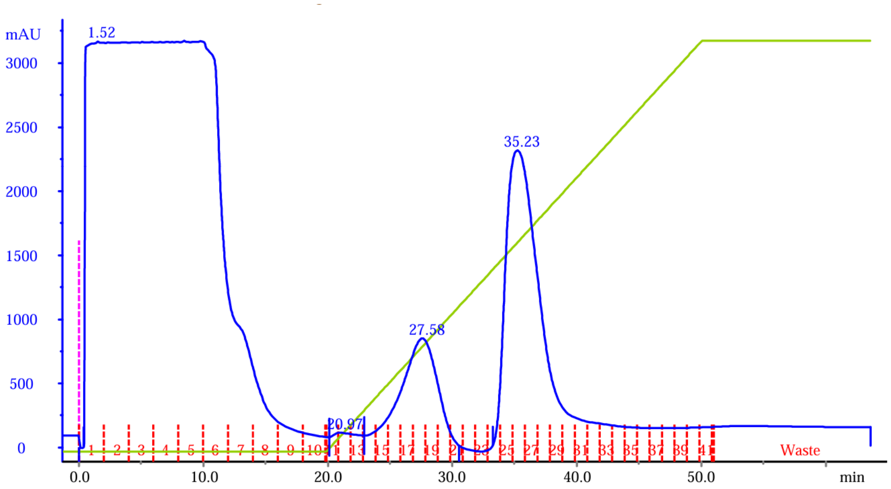


**Figure S16. Representative FPLC trace from MrtR purification.** Blue line represents absorbance at 280 nm. Green line represents percent of buffer B, which contains 360 mM imidazole, from 0 to 100%. Red lines indicate fractions collected. Peaks 1 and 2 (at 27.58 and 35.23 min, respectively) both contained MrtR.

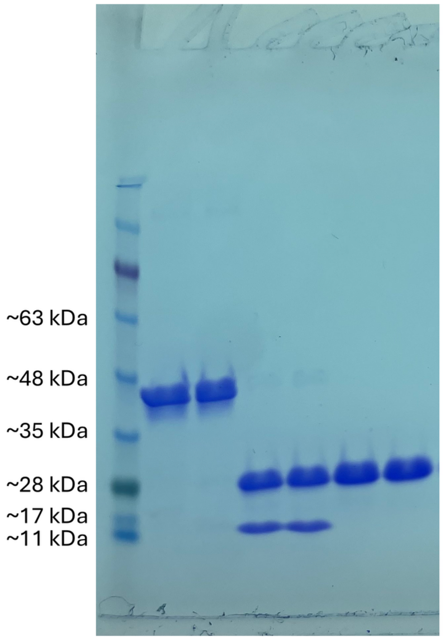


**Figure S17. Representative SDS-PAGE gel from MrtR purification.** From left to right, lanes represent protein ladder → peak 1 (from FPLC trace in **Figure S16**) → peak 2 → peak 1 after His6-SUMO tag cleavage → peak 2 after tag cleavage → peak 1 after tag removal → peak 2 after tag removal.

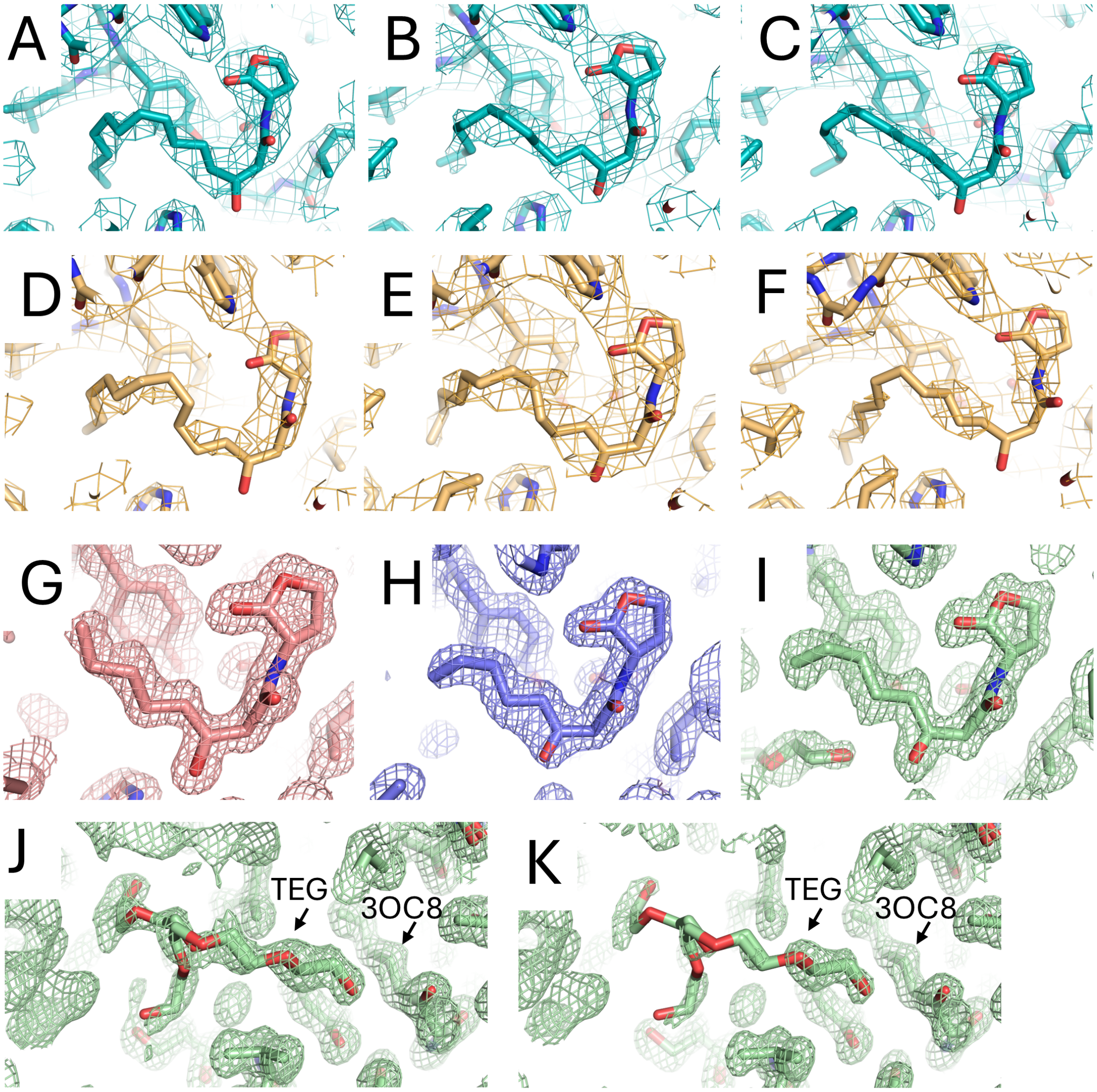


**Figure S18. Omit maps showing electron density calculated without the AHL modeled.** Models with ligand are shown with omit maps to indicate ligand position. Images for MrtR:7*Z*-3OHC14 (**A**–**C**), MrtR:3OHC14 (**D**–**F**), MrtR:3OHC8 (**G**), MrtR:3OC8 (**H**), and MrtR:3OC8+TEG (**I**) structures. Omit maps showing electron density for MrtR:3OC8+TEG without the tetraethylene glycol (TEG) or AHL modeled at 0.7σ (**J**) and 1σ (**K**).

**Table S1. Abbreviated table of BLAST results for MrtR.** Hits were selected to reflect the breadth of species and strains from a BLAST (26) search that provided 500 results. Query cover was 100% for all results. Per. Ident. indicates percent identity, and Acc. Len. indicates accession length.

| Description | Scientific Name | E value | Per. Ident. | Acc. Len | NCBI accession |
|---|---|---|---|---|---|
| MULTISPECIES: LuxR family transcriptional regulator | *Rhizobium* | 1.00E-177 | 100 | 241 | WP_017961376.1 |
| LuxR family transcriptional regulator | *Rhizobium leguminosarum* | 2.00E-177 | 99.6 | 241 | WP_027664189.1 |
| LuxR family transcriptional regulator | *Rhizobium laguerreae* | 6.00E-177 | 99.2 | 241 | WP_168284029.1 |
| LuxR family transcriptional regulator | *Rhizobium aouanii* | 2.00E-176 | 99.2 | 241 | WP_335910586.1 |
| LuxR family transcriptional regulator | *Rhizobium ruizarguesonis* | 4.00E-176 | 98.3 | 241 | WP_018074074.1 |
| LuxR family transcriptional regulator | *Rhizobium leguminosarum* | 5.00E-176 | 98.3 | 241 | WP_129419792.1 |
| LuxR family transcriptional regulator | *Rhizobium changzhiense* | 5.00E-176 | 98.8 | 241 | WP_180695424.1 |
| LuxR transcriptional regulator CinR | *Rhizobium leguminosarum* bv. *viciae* | 2.00E-175 | 98.3 | 243 | AVC50771.1 |
| LuxR family transcriptional regulator | *Rhizobium sophorae* | 2.00E-175 | 98.3 | 241 | WP_168309920.1 |
| LuxR family transcriptional regulator | *Rhizobium indigoferae* | 1.00E-174 | 97.9 | 241 | WP_193445686.1 |
| LuxR family transcriptional regulator | *Rhizobium phaseoli* | 3.00E-174 | 97.1 | 241 | WP_164011839.1 |
| LuxR family transcriptional regulator | *Rhizobium esperanzae* | 3.00E-174 | 97.5 | 241 | WP_088392013.1 |
| LuxR family transcriptional regulator | *Rhizobium lentis* | 1.00E-173 | 97.1 | 241 | WP_183912585.1 |
| LuxR family transcriptional regulator | *Rhizobium etli* | 2.00E-173 | 96.7 | 241 | WP_020921838.1 |

| | | | | | |
|---|---|---|---|---|---|
| autoinducer transcriptional regulator protein | *Rhizobium etli* CFN 42 | 9.00E-173 | 96.7 | 260 | ABC91683.1 |
| LuxR family transcriptional regulator | *Rhizobium sophoriradicis* | 2.00E-172 | 96.7 | 241 | WP_096770464.1 |
| LuxR family transcriptional regulator | *Rhizobium leguminosarum* | 3.00E-172 | 95.9 | 241 | WP_116406551.1 |
| LuxR family transcriptional regulator | *Rhizobium lentis* | 3.00E-172 | 96.3 | 241 | WP_221126963.1 |
| LuxR family transcriptional regulator | *Rhizobium bangladeshense* | 3.00E-172 | 96.7 | 241 | WP_064684468.1 |
| CinR | *Rhizobium etli* | 6.00E-172 | 96.3 | 241 | AAL59595.1 |

**Table S2. Abbreviated table of BLAST results for the promoter sequence in the *E. coli* MrtR transcriptional reporter.** Hits were selected to reflect the breadth of species and strains from a BLAST (26) search that provided 100 results. Per. Ident. indicates percent identity, and Acc. Len. indicates accession length.

| Description | Scientific Name | Query Cover | E value | Per. Ident. | NCBI accession |
|---|---|---|---|---|---|
| *Rhizobium* sp. SU303 chromosome, complete genome | *Rhizobium* sp. SU303 | 100% | 8.00E-54 | 100 | CP088095.1 |
| *Rhizobium laguerreae* strain WSM1455 chromosome, complete genome | *Rhizobium laguerreae* | 100% | 8.00E-54 | 100 | CP088090.1 |
| *Rhizobium leguminosarum* bv. *viciae* strain BIHB 1148 plasmid pSK03, complete sequence | *Rhizobium leguminosarum* bv. *viciae* | 100% | 8.00E-54 | 100 | CP022567.1 |
| *Rhizobium brockwellii* strain CC275e chromosome, complete genome | *Rhizobium brockwellii* | 100% | 5.00E-46 | 95.9 | CP053439.1 |
| *Rhizobium ruizarguesonis* strain TP15 chromosome, complete genome | *Rhizobium ruizarguesonis* | 100% | 5.00E-41 | 93.5 | CP140868.1 |
| *Rhizobium indicum* strain JKLM 13E chromosome, complete genome | *Rhizobium indicum* | 100% | 5.00E-41 | 93.5 | CP054031.1 |
| *Rhizobium johnstonii* strain 3841 chromosome, complete genome | *Rhizobium johnstonii* | 98% | 6.00E-40 | 93.4 | CP176004.1 |
| *Rhizobium anhuiense* bv. *trifolii* strain T24 chromosome, complete genome | *Rhizobium anhuiense* bv. *trifolii* | 81% | 5.00E-36 | 96.9 | CP098803.1 |
| *Rhizobium indigoferae* strain CIP 108029 chromosome, complete genome | *Rhizobium indigoferae* | 100% | 6.00E-35 | 90.3 | CP140635.1 |
| *Rhizobium leguminosarum* strain ATCC 14479 chromosome, complete genome | *Rhizobium leguminosarum* | 79% | 1.00E-31 | 94.7 | CP030760.1 |

| | | | | | |
|---|---|---|---|---|---|
| *Rhizobium hidalgonense* strain JKLM 19E chromosome, complete genome | *Rhizobium hidalgonense* | 78% | 5.00E-31 | 94.7 | CP054027.1 |
| *Rhizobium lentis* strain BLR27 chromosome, complete genome | *Rhizobium lentis* | 78% | 5.00E-31 | 94.7 | CP071454.1 |
| *Rhizobium leguminosarum* bv. *trifolii* WSM2304 chromosome, complete genome | *Rhizobium leguminosarum* bv. *trifolii* WSM2304 | 78% | 1.00E-27 | 92.6 | CP001191.1 |
| *Rhizobium etli* CFN 42, complete genome | *Rhizobium etli* CFN 42 | 79% | 1.00E-27 | 92.6 | CP000133.1 |
| *Rhizobium bangladeshense* strain PLR8-1a chromosome, complete genome | *Rhizobium bangladeshense* | 79% | 5.00E-26 | 91.6 | CP071618.1 |

**Table S3. Origins of compounds tested in this study.** Naturally occurring AHLs acquired from commercial sources are excluded.

<table>
<tr><th>Compound</th><th>References</th></tr>
<tr><td>B7</td><td rowspan="7">(27)</td></tr>
<tr><td>B10</td></tr>
<tr><td>D15</td></tr>
<tr><td>C6</td></tr>
<tr><td>C10</td></tr>
<tr><td>C11</td></tr>
<tr><td>C14</td></tr>
<tr><td>K3</td><td>(28, 29)</td></tr>
<tr><td>E33</td><td>(30)</td></tr>
<tr><td>H23</td><td>(31, 32)</td></tr>
<tr><td>V-06-018</td><td>(33, 34)</td></tr>
<tr><td>TP-5</td><td>(34, 35)</td></tr>
<tr><td>CL</td><td>(34, 36)</td></tr>
<tr><td>S1</td><td rowspan="3">(37)</td></tr>
<tr><td>S2</td></tr>
<tr><td>S3</td></tr>
<tr><td>3R-OHC12</td><td rowspan="2">(38)</td></tr>
<tr><td>3S-OHC12</td></tr>
<tr><td>3OHC8</td><td>(32)</td></tr>
<tr><td>3OC16</td><td>(32); See supplemental methods.</td></tr>
<tr><td>3OC18</td><td rowspan="2">See supplemental methods.</td></tr>
<tr><td>3OC20</td></tr>
</table>

**Table S4. Molecular weight of MrtR complexes estimated based on size exclusion chromatography.** Values are the average of two replicate runs. Full traces for samples and calibration standards shown at the end of this SI Appendix.

| Condition | Molecular weight (kDa) (standard deviation) |
|---|---|
| DMSO | 25.9 (0.1) |
| 3OC8 | 26.9 (0.3) |
| 3OC12 | 26.8 (0.2) |
| 7*Z*-3OHC14 | 48.1 (0.0) |
| C14-HSL | 42.7 (0.1) |

**Table S5. X-ray crystallographic data collection and refinement statistics.**[a]

| PDB ID | 9Y2H | 9ZZ0 | 9ZPJ | 11BM | 10ZI |
|---|---|---|---|---|---|
| Ligand | 7Z-3OHC14 | 3OHC14 | 3OHC8 | 3OC8 | 3OC8 + TEG |
| **Data Collection** | | | | | |
| Space group | C121 | C121 | C121 | C121 | C121 |
| Cell dimensions (Å) | a, b, c = 67.10, 120.79, 122.60 | a, b, c = 67.04,120.93, 122.54 | a, b, c = 125.32, 35.55, 61.65 | a, b, c = 126.33, 35.55, 61.96 | a, b, c = 126.43, 35.73, 61.94 |
| Cell angles | a = γ = 90.00, β = 98.58 | α = γ = 90.00, β = 98.81 | α = γ = 90.00, β = 111.66 | α = γ = 90.00, β = 112.51 | α = γ = 90.00, β = 112.13 |
| Wavelength (Å) | 0.97934 | 0.97934 | 0.98011 | 0.96546 | 0.96546 |
| Beamline | NSLS-II 17-ID-2 | NSLS-II 17-ID-2 | ESRF BM07 | ESRF MASSIF-1 | ESRF MASSIF-1 |
| Resolution (Å) | 42.8 – 3.06 (3.11 – 3.06) | 34.4-3.36 (3.42-3.36) | 51.4-1.18 (1.20-1.18) | 57.2-1.44 (1.47-1.44) | 58.6-1.22 (1.24-1.22) |
| Unique reflections | 17,089 (909) | 12,733 (519) | 77,750 (4011) | 46,325 (2324) | 70,635 (3460) |
| Completeness (%) | 93.2 (100) | 92.2 (76.2) | 92.4 (96.4) | 100 (100) | 92.6 (90.5) |
| $CC_{1/2}$ | 0.954 (0.350) | 0.930 (0.324) | 0.999 (0.385) | 0.998 (0.402) | 0.999 (0.681) |
| I/σ(I) | 4.0 (1.2) | 4.2 (1.4) | 13.5 (0.8) | 10.8 (0.6) | 11.5 (0.5) |
| Redundancy | 6.1 (6.4) | 6.9 (6.5) | 6.5 (6.3) | 6.7 (5.8) | 6.4 (4.5) |
| **Refinement** | | | | | |
| Total no. of reflections | 16,275 | 12,137 | 73,709 | 43,609 | 66,060 |
| Total no. of non-hydrogen atoms | 5,591 | 5,437 | 2613 | 2,121 | 2,751 |
| Final bin resolution (Å) | (3.14 – 3.06) | (3.45-3.36) | (1.21-1.18) | (1.48-1.44) | (1.25-1.22) |
| $R_{work}$ (%) | 25.6 (37.8) | 25.3 (32.9) | 14.6 (32.3) | 18.2 (37.2) | 15.5 (42.3) |
| $R_{free}$ test set (%) | 4.7 | 4.7 | 5.0 | 5.0 | 5.0 |
| $R_{free}$ (%) | 29.4 (37.3) | 30.8 (34.9) | 17.3 (32.6) | 23.2 (36.6) | 19.3 (43.5) |
| Average B factor ($Å^2$) | 62.3 | 56.7 | 21.4 | 30.2 | 18.3 |
| Ramachandran plot Favored (%) | 97.9 | 97.3 | 99.2 | 100 | 99.6 |
| Outliers (%) | 0 | 0 | 0 | 0 | 0 |

[a] Values in parenthesis are for the highest resolution shell. $R_{work} = \Sigma||F_o - F_c|| / F_o$, where $F_o$ is an observed amplitude and $F_c$ a calculated amplitude. $R_{free}$ is the same statistic calculated with a subset of the data that was excluded from refinement. Ramachandran statistics were calculated with Molprobity (http://molprobity.biochem.duke.edu/) (39).

**Table S6. Assemblies predicted for MrtR:7*Z*-3OHC14 by PDBePISA** (15)**.** No higher-order assemblies were predicted.[a]

| Composition | Id | Stable | Surface area ($Å^2$) | Buried area ($Å^2$) | $ΔG^{int}$ (kcal/mol) | $ΔG^{diss}$ (kcal/mol) |
|---|---|---|---|---|---|---|
| $A_2[LIG]_2$ | 1 | yes | 20880 | 5410 | -22.4 | 12.9 |
| $BC[LIG]_2$ | 1 | yes | 20580 | 5490 | -20.9 | 11.6 |

[a] **Composition** indicates monomeric units found in the assembly, with A, B, and C representing the chains in the asymmetric unit. **Id** indicates distinctive assembly types, with structurally identical assemblies assigned the same Id. **Stable** indicates whether the assembly is predicted to be stable in solution. **Surface area** indicates the solvent-accessible surface area of the assembly. **Buried area** indicates the solvent-accessible surface area of monomeric units buried upon assembly formation. **$ΔG^{int}$** indicates the solvation free energy gain upon formation of the assembly. The value is calculated as the difference in total solvation energies between isolated and assembled structures. This value does not include the effect of satisfied hydrogen bonds and salt bridges across the assembly's interfaces. **$ΔG^{diss}$** indicates the free energy difference between dissociated and associated states. Assemblies with **$ΔG^{diss} > 0$** are thermodynamically stable.

**Table S7. Assemblies predicted for MrtR:3OHC14 by PDBePISA** (15)**.** No higher-order assemblies were predicted. Assemblies are analogous to those predicted for MrtR:7*Z*-3OHC14.[a]

| Composition | Id | Stable | Surface area ($Å^2$) | Buried area ($Å^2$) | $ΔG^{int}$ (kcal/mol) | $ΔG^{diss}$ (kcal/mol) |
|---|---|---|---|---|---|---|
| $C_2[LIG]_2$ | 1 | yes | 20470 | 5120 | -23.8 | 15.1 |
| $AB[LIG]_2$ | 1 | yes | 20340 | 5210 | -17.1 | 10.4 |

[a] See **Table S6** footnote for term definitions.

**Table S8. Bacterial strains and plasmids used in this study.**

| Strain | Description | Reference |
|---|---|---|
| *E. coli* Top 10 | Used for cloning. | Lab collection |
| *E. coli* BW27749 | Constitutive expression of low-affinity arabinose pump AraE and deletion of *araFGH*. Used for reporters. | (40) |
| *E. coli* BL21(DE3) | Used for protein overexpression. | Novagen |
| **Plasmid** | **Description** | **Reference** |
| pJN105-mrtR | pJN105 with arabinose-inducible promoter controlling *mrtR* ($Gm^R$). | (2) |
| pmrtI:GFP | *mrtI* promoter (-120 to -1 relative the *mrtI* translational start site) from *Mesorhizobium tianshanense* controlling *GFP* ($Ap^R$). | (2) |
| pJN105-mrtR R57M | pJN105-mrtR with an R57M mutation. | This study |
| pJN105-mrtR S72A | pJN105-mrtR with an S72A mutation. | This study |
| pJN105-mrtR Q45M | pJN105-mrtR with a Q45M mutation. | This study |
| pJN105-mrtR T46V | pJN105-mrtR with a T46V mutation. | This study |
| pJN105-mrtR D52L | pJN105-mrtR with a D52L mutation. | This study |
| pJN105-mrtR Q45M, R57M | pJN105-mrtR with Q45M and R57M mutations. | This study |
| pJN105-mrtR R57M, S72A | pJN105-mrtR with R57M and S72A mutations. | This study |
| pJN105-mrtR Q45M, S72A | pJN105-mrtR with Q45M and S72A mutations. | This study |
| pJN105-mrtR D174A | pJN105-mrtR with a D174A mutation. | This study |
| pJN105-mrtR I47A | pJN105-mrtR with an I47A mutation. | This study |
| pJN105-mrtR V176A | pJN105-mrtR with a V176A mutation. | This study |
| pJN105-mrtR L236A | pJN105-mrtR with an L236A mutation. | This study |
| pJN105-mrtR I47A, L236A | pJN105-mrtR with I47A and L236A mutations. | This study |
| pJN105-mrtR V176A, L236A | pJN105-mrtR with V176A and L236A mutations | This study |
| pJN105-mrtR I47A, V176A, L236A | pJN105-mrtR with I47A, V176A, and L236A mutations. | This study |
| pJN105-mrtR L69A | pJN105-mrtR with an L69A mutation. | This study |
| pJN105-mrtR L69F | pJN105-mrtR with an L69F mutation. | This study |
| pJN105-mrtR H42L | pJN105-mrtR with an H42L mutation. | This study |
| pJN105-mrtR V74A | pJN105-mrtR with a V74 mutation. | This study |
| pJN105-mrtR P54A | pJN105-mrtR with a P54A mutation. | This study |
| pJN105-mrtR V65A | pJN105-mrtR with a V65A mutation. | This study |
| pJN105-mrtR L70A | pJN105-mrtR with an L70A mutation. | This study |
| pJN105-mrtR S66A | pJN105-mrtR with an S66A mutation. | This study |

| | | |
|---|---|---|
| pJN105-mrtR D62L | pJN105-mrtR with a D52L mutation. | This study |
| pJN105-mrtR I51A | pJN105-mrtR with an I51A mutation. | This study |
| pJN105-mrtR S66A, D62L | pJN105-mrtR with S66A and D62L mutations. | This study |
| pJN105-mrtR V56A | pJN105-mrtR with a V56A mutation. | This study |
| pJN105-mrtR S66A, D62L, V65A, L70A | pJN105-mrtR with S66A, D52L, V65A, and L70A mutations. | This study |
| pET28-his6-SUMO-smaR | Used to amplify vector for MrtR overexpression vector. | GenScript |
| pET28-his6-SUMO-mrtR | Vector for overexpression of MrtR. | This study |

**Table S9. Primers and oligos used in this study.**

| Name | Sequence |
|---|---|
| Nonspecific competitor for FP (pUC18-MCS)_fwd | tacggacgtccagctgagatctcctaggggcc |
| Nonspecific competitor for FP (pUC18-MCS)_rev | ggcccctaggagatctcagctggacgtccgta |
| FAM-labeled MrtR binding site for FP_fwd | /56-FAM/ttgcgatatgcgcCCCCCTCATCTGAGGGGGcccatctgagg |
| Specific competitor MrtR binding site for FP_fwd | ttgcgatatgcgcCCCCCTCATCTGAGGGGGcccatctgagg |
| MrtR binding site for FP_rev, used with FAM and specific competitor fwd strands | cctcagatgggCCCCCTCAGATGAGGGGGgcgcatatcgcaa |
| MrtR Q45M_fwd | tcacttggcgatgaccatcgccg |
| MrtR Q45M_rev | tacgtgacgaaatccaacc |
| MrtR L69A_fwd | cagccgctacgcgctgaacagct |
| MrtR L69A_rev | acccacgcatcagga |
| MrtR L69F_fwd | cagccgctactttctgaacagct |
| MrtR L69F_rev | acccacgcatcaggatat |
| MrtR S72A_fwd | cctgctgaacgcgtatgtcaaggtggacc |
| MrtR S72A_rev | tagcggctgacccac |
| MrtR R57M_fwd | gccattcgttatgactacatatcctgatgc |
| MrtR R57M_rev | gagtcgattttggcg |
| MrtR D52L_fwd | cgccaaaatcctgtcgccattcgttcgc |
| MrtR D52L_rev | gcgatggtttgcgcc |
| MrtR T46V_fwd | cttggcgcaagtgatcgccgcca |
| MrtR T46V_rev | tgatacgtgacgaaatccaac |
| MrtR H42L_fwd | cgtcacgtatctgttggcgcaaac |
| MrtR H42L_rev | aaatccaacccatattctg |

| | |
|---|---|
| MrtR I47A_fwd | ggcgcaaaccgcggccgccaaaatc |
| MrtR I47A_rev | aagtgatacgtgacgaaatc |
| MrtR L236A_fwd | tgccgttcaggcgcgcatcatcaatccgtaaatgatc |
| MrtR L236A_rev | cggctggccgctgct |
| MrtR V176A_fwd | gaacgatccggcgcccgcccttt |
| MrtR V176A_rev | tccccgtgcaactcg |
| MrtR V74A_fwd | gaacagctatgcgaaggtggacccaatcg |
| MrtR V74A_rev | agcaggtagcggctgacc |
| MrtR P54A_fwd | aatcgactcggcgttcgttcgcac |
| MrtR P54A_rev | ttggcggcgatggtttgc |
| MrtR V65A_fwd | tgatgcgtgggcgagccgctacc |
| MrtR V65A_rev | ggatatgtagtgcgaacgaatgg |
| MrtR L70A_fwd | ccgctacctggcgaacagctatgtc |
| MrtR L70A_rev | ctgacccacgcatcagga |
| MrtR S66A_fwd | tgcgtgggtcgcgcgctacctgc |
| MrtR S66A_rev | tcaggatatgtagtgcgaac |
| MrtR D62L_fwd | tacatatcctctggcgtgggtcagc |
| MrtR D62L_rev | gtgcgaacgaatggcgag |
| MrtR I51A_fwd | cgccgccaaagcggactcgccattc |
| MrtR I51A_rev | atggtttgcgccaagtga |
| MrtR V56A_fwd | ctcgccattcgcgcgcactacatatc |
| MrtR V56A_rev | tcgattttggcggcgatg |
| MrtR S66A, D62L, V65A, L70A_fwd | cgcgctacctggcgaacagctatgtcaaggtg |
| MrtR S66A, D62L, V65A, L70A_rev | ccgcccacgccagaggatatgtagtgcgaac |
| pET-his6-SUMO-mrtR vector_fwd | ttcgcatcatcaatccgtaactcgagctcgagcaccacca |
| pET-his6-SUMO-mrtR vector_rev | gaatatgtgttttcgatcataccaccaatctgttctctgt |
| pET-his6-SUMO-mrtR insert_fwd | acagagaacagattggtggtatgatcgaaaacacatattcgg |
| pET-his6-SUMO-mrtR insert_rev | tggtggtgctcgagctcgagttacggattgatgatgcgaa |

### Relative fluorescence unit (RFU) traces for MrtR DSF.

Traces correspond to data shown in **Figure 1D** and **Table 1** and represent three runs of three technical replicates, except for the buffer only condition, which represents one run. Compound names are indicated at the top of each trace.

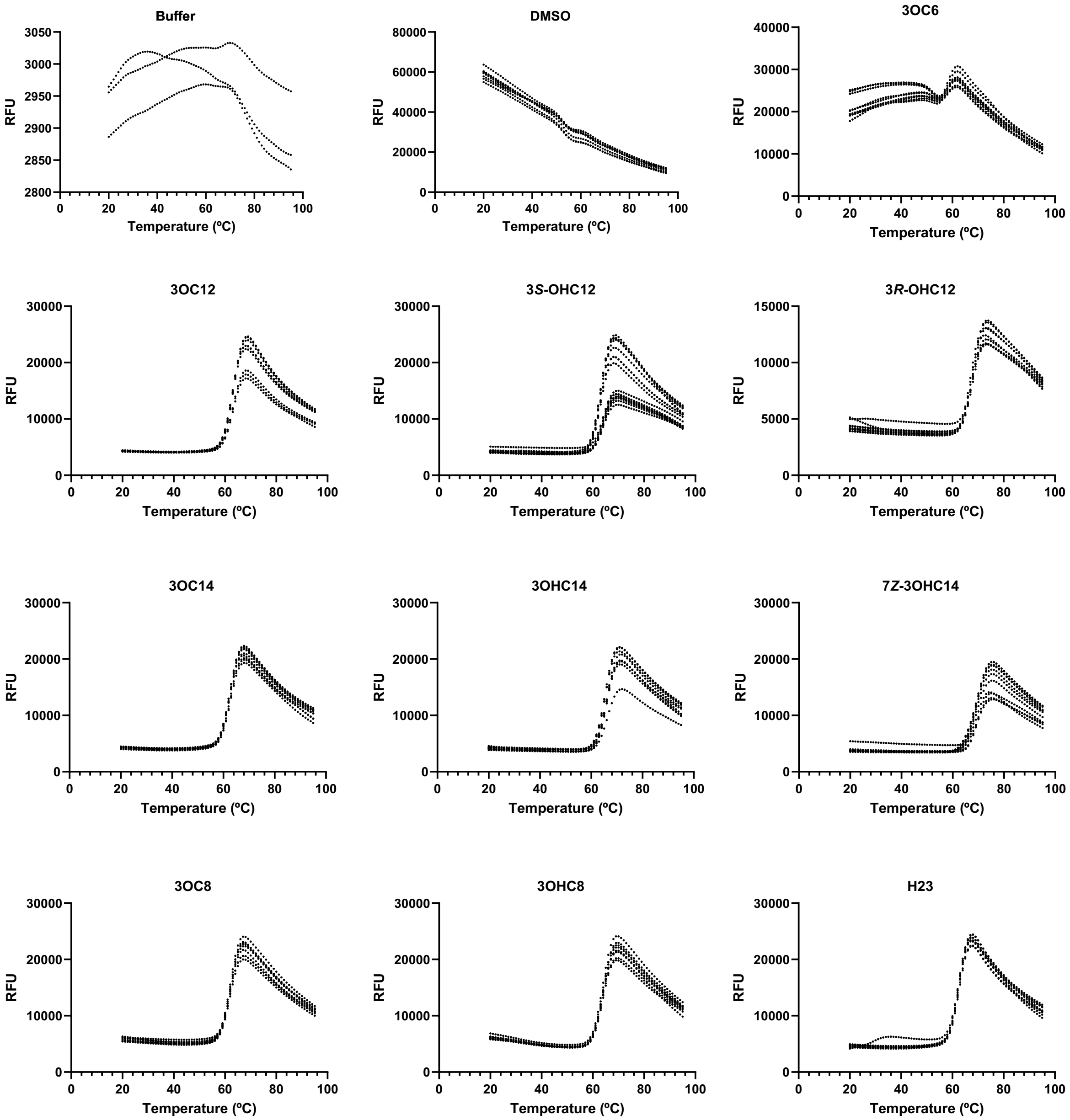

**<u>First derivative traces for MrtR DSF.</u>**

Traces correspond to data shown in **Figure 1D** and **Table 1** and represent three runs of three technical replicates, except for the buffer only condition, which represents one run. Compound names are indicated at the top of each trace.

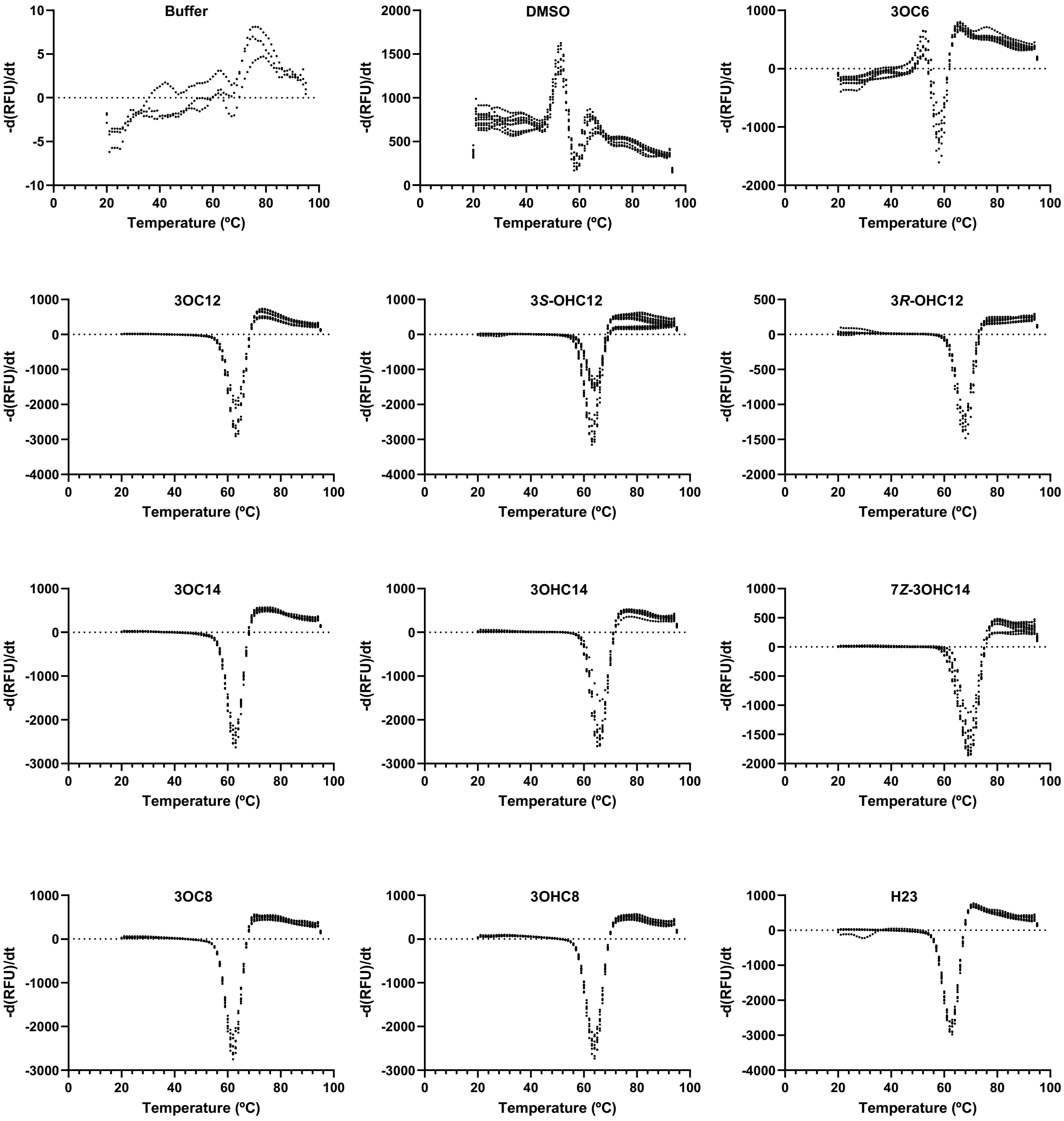

**<u>Size exclusion chromatography traces used to estimate MrtR complex molecular weight.</u>**

Data are summarized in **Figure 1E** and **Table S4**. Sample names are indicated at the top of each trace.

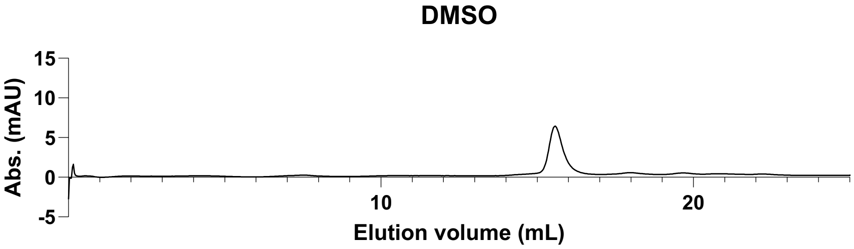


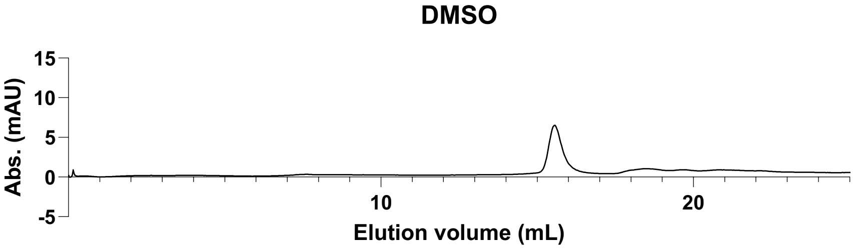


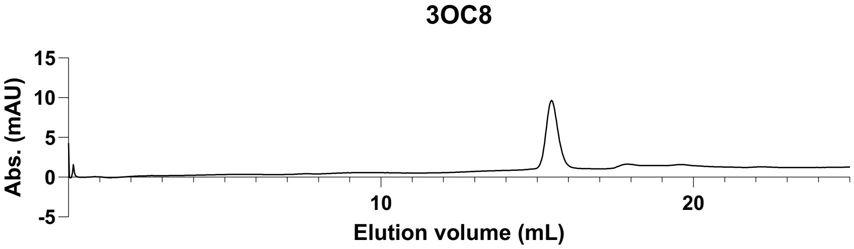


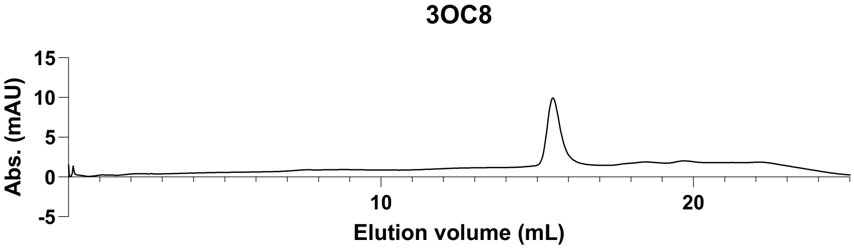


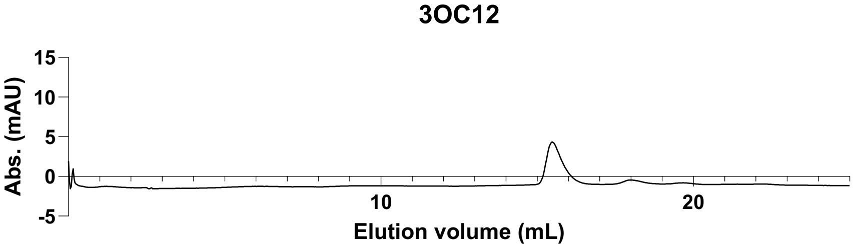

**3OC12**

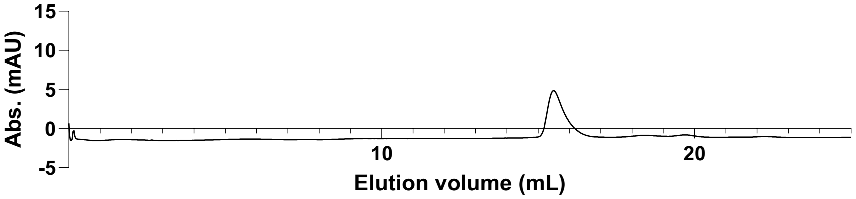


**C14-HSL**

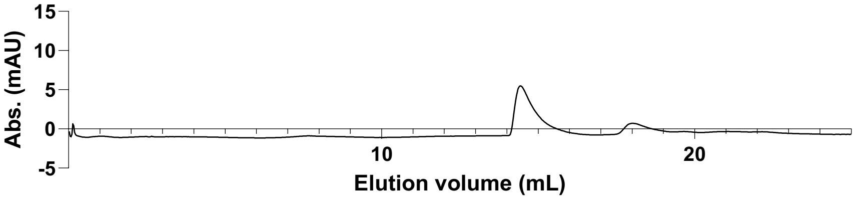


**C14-HSL**

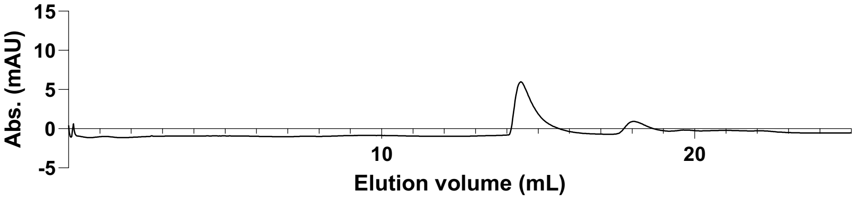


**7Z-3OH14**

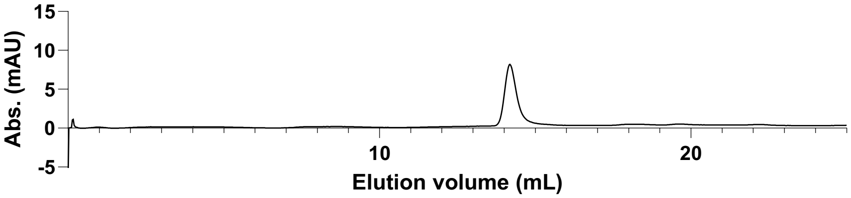


**7Z-3OH14**

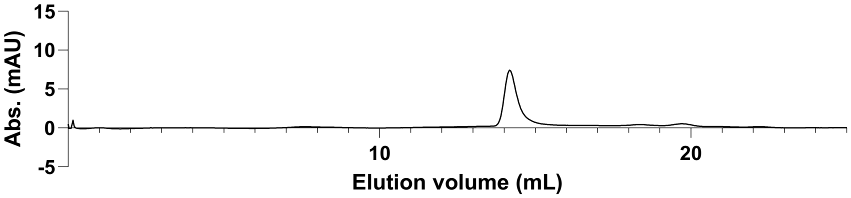

**Standards with DMSO**

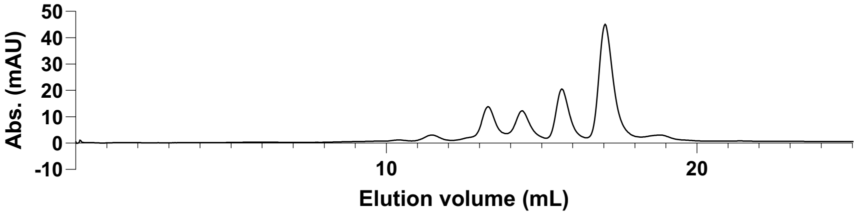


**Standards with 3OC12**

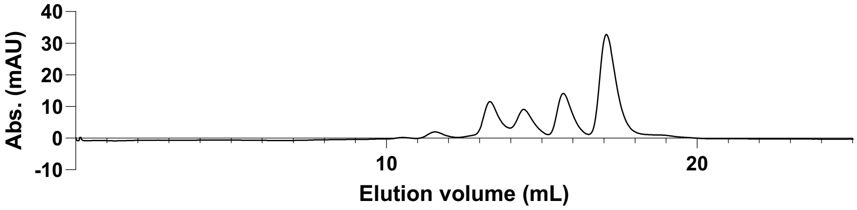


**Standards with 7*Z*-3OHC14**

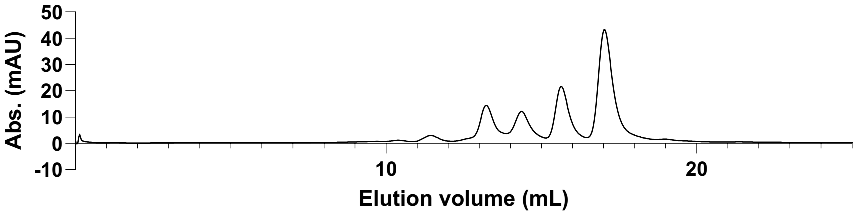


**No protein injected**

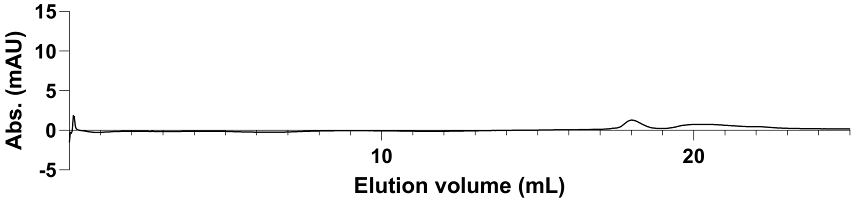


**Running buffer injected**

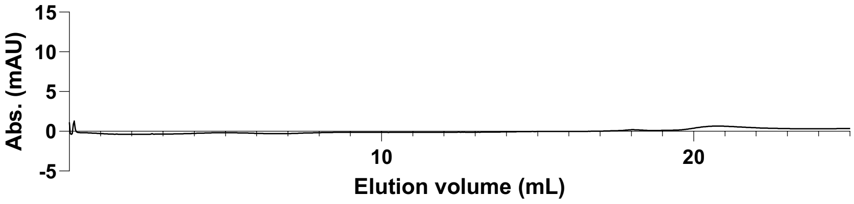

**References.**
