## Supplementary material for "MrtR of *Mesorhizobium tianshanense* reveals both activation and inhibition mechanisms of a LuxR-type quorum sensing receptor": Structure validation reports: MrtR_3OC8_11BM.pdf

### Full wwPDB X-ray Structure Validation Report ⓘ

Feb 17, 2026 – 07:04 PM EST

| | | |
|---|---|---|
| PDB ID | : | 11BM / pdb_000011bm |
| Title | : | Crystal structure of MrtR bound to 3O-C8 homoserine lactone |
| Deposited on | : | 2026-02-16 |
| Resolution | : | 1.44 Å(reported) |

**This wwPDB validation report is for manuscript review**

This is a Full wwPDB X-ray Structure Validation Report.

This report is produced by the wwPDB biocuration pipeline after annotation of the structure.

We welcome your comments at **
A user guide is available at
https://www.wwpdb.org/validation/2017/XrayValidationReportHelp
with specific help available everywhere you see the ⓘ symbol.

The types of validation reports are described at
http://www.wwpdb.org/validation/2017/FAQs#types.

For Manuscript Review

The following versions of software and data (see references ⓘ) were used in the production of this report:

| | | |
|---|---|---|
| MolProbity | : | 4-5-2 with Phenix2.0 |
| Mogul | : | 2022.3.0, CSD as543be (2022) |
| Xtriage (Phenix) | : | 2.0 |
| EDS | : | 3.0 |
| buster-report | : | 1.1.7 (2018) |
| Percentile statistics | : | 20231227.v01 (using entries in the PDB archive December 27th 2023) |
| CCP4 | : | 9.0.010 (Gargrove) |
| Density-Fitness | : | 1.0.12 |
| Ideal geometry (proteins) | : | Engh & Huber (2001) |
| Ideal geometry (DNA, RNA) | : | Parkinson et al. (1996) |

### 1 Overall quality at a glance (i)

The following experimental techniques were used to determine the structure:
*X-RAY DIFFRACTION*

The reported resolution of this entry is 1.44 Å.

Percentile scores (ranging between 0-100) for global validation metrics of the entry are shown in the following graphic. The table shows the number of entries on which the scores are based.

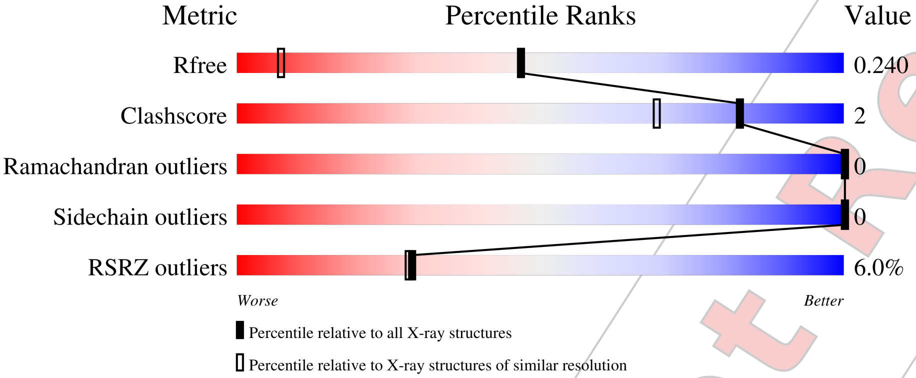


| Metric | Whole archive (#Entries) | Similar resolution (#Entries, resolution range(Å)) |
|---|---|---|
| $R_{free}$ | 164625 | 2809 (1.46-1.42) |
| Clashscore | 180529 | 3008 (1.46-1.42) |
| Ramachandran outliers | 177936 | 2971 (1.46-1.42) |
| Sidechain outliers | 177891 | 2971 (1.46-1.42) |
| RSRZ outliers | 164620 | 2809 (1.46-1.42) |

The table below summarises the geometric issues observed across the polymeric chains and their fit to the electron density. The red, orange, yellow and green segments of the lower bar indicate the fraction of residues that contain outliers for >=3, 2, 1 and 0 types of geometric quality criteria respectively. A grey segment represents the fraction of residues that are not modelled. The numeric value for each fraction is indicated below the corresponding segment, with a dot representing fractions <=5% The upper red bar (where present) indicates the fraction of residues that have poor fit to the electron density. The numeric value is given above the bar.

| Mol | Chain | Length | Quality of chain |
|---|---|---|---|
| 1 | A | 241 | 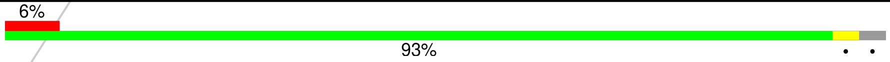 |



For Manuscript Review

#### 2 Entry composition (i)

There are 4 unique types of molecules in this entry. The entry contains 2121 atoms, of which 0 are hydrogens and 0 are deuteriums.

In the tables below, the ZeroOcc column contains the number of atoms modelled with zero occupancy, the AltConf column contains the number of residues with at least one atom in alternate conformation and the Trace column contains the number of residues modelled with at most 2 atoms.

- Molecule 1 is a protein called MrtR.

| Mol | Chain | Residues | Atoms | | | | | ZeroOcc | AltConf | Trace |
|---|---|---|---|---|---|---|---|---|---|---|
| | | | Total | C | N | O | S | | | |
| 1 | A | 233 | 1894 | 1209 | 330 | 350 | 5 | 0 | 7 | 0 |

- Molecule 2 is 3-OXO-OCTANOIC ACID (2-OXO-TETRAHYDRO-FURAN-3-YL)-AMIDE (CCD ID: LAE) (formula: $C_{12}H_{19}NO_4$) (labeled as "Ligand of Interest" by depositor).

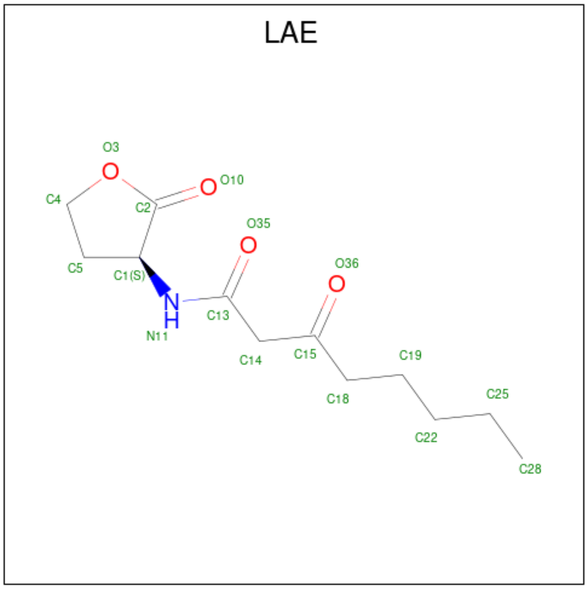


| Mol | Chain | Residues | Atoms | | | | ZeroOcc | AltConf |
|---|---|---|---|---|---|---|---|---|
| | | | Total | C | N | O | | |
| 2 | A | 1 | 17 | 12 | 1 | 4 | 0 | 0 |

- Molecule 3 is DI(HYDROXYETHYL)ETHER (CCD ID: PEG) (formula: $C_4H_{10}O_3$).

For Manuscript Review

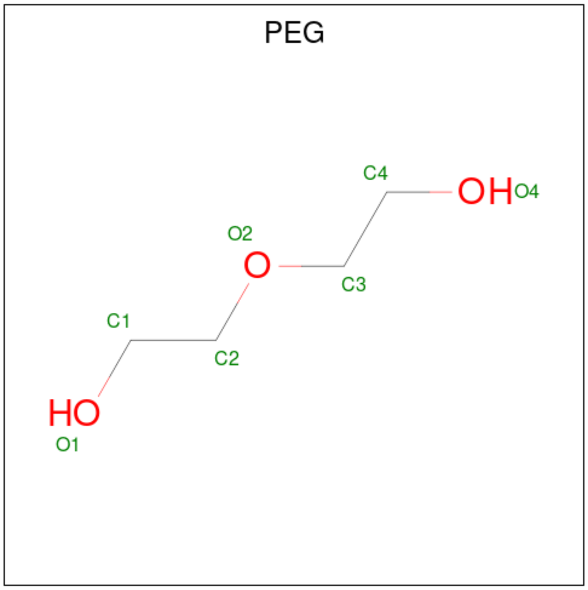


| Mol | Chain | Residues | Atoms | ZeroOcc | AltConf |
|---|---|---|---|---|---|
| 3 | A | 1 | Total C O<br>7 4 3 | 0 | 0 |
| 3 | A | 1 | Total C O<br>7 4 3 | 0 | 0 |

- Molecule 4 is water.

| Mol | Chain | Residues | Atoms | ZeroOcc | AltConf |
|---|---|---|---|---|---|
| 4 | A | 196 | Total O<br>196 196 | 0 | 0 |

For Manuscript Review

### 3 Residue-property plots (i)

These plots are drawn for all protein, RNA, DNA and oligosaccharide chains in the entry. The first graphic for a chain summarises the proportions of the various outlier classes displayed in the second graphic. The second graphic shows the sequence view annotated by issues in geometry and electron density. Residues are color-coded according to the number of geometric quality criteria for which they contain at least one outlier: green = 0, yellow = 1, orange = 2 and red = 3 or more. A red dot above a residue indicates a poor fit to the electron density (RSRZ > 2). Stretches of 2 or more consecutive residues without any outlier are shown as a green connector. Residues present in the sample, but not in the model, are shown in grey.

• Molecule 1: MrtR

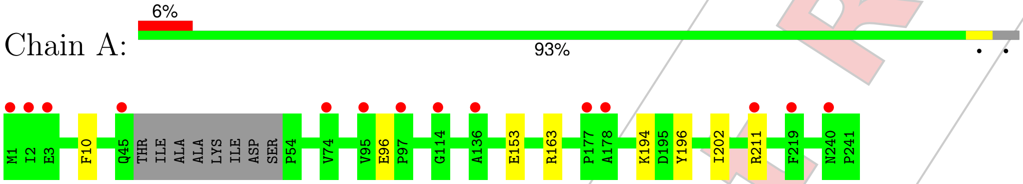


For Manuscript Review

### 4 Data and refinement statistics (i)

| Property | Value | Source |
|---|---|---|
| Space group | C 1 2 1 | Depositor |
| Cell constants<br>a, b, c, $\alpha$, $\beta$, $\gamma$ | 126.33Å 35.55Å 61.96Å<br>90.00° 112.51° 90.00° | Depositor |
| Resolution (Å) | 57.20 – 1.44<br>57.20 – 1.44 | Depositor<br>EDS |
| % Data completeness<br>(in resolution range) | 99.2 (57.20-1.44)<br>99.2 (57.20-1.44) | Depositor<br>EDS |
| $R_{merge}$ | (Not available) | Depositor |
| $R_{sym}$ | (Not available) | Depositor |
| $< I/\sigma(I) >$ [1] | 1.37 (at 1.44Å) | Xtriage |
| Refinement program | REFMAC 5.8.0430 | Depositor |
| R, $R_{free}$ | 0.182 , 0.232<br>0.191 , 0.240 | Depositor<br>DCC |
| $R_{free}$ test set | 2318 reflections (5.01%) | wwPDB-VP |
| Wilson B-factor (Å$^2$) | 23.3 | Xtriage |
| Anisotropy | 0.415 | Xtriage |
| Bulk solvent $k_{sol}$(e/Å$^3$), $B_{sol}$(Å$^2$) | 0.32 , 44.2 | EDS |
| L-test for twinning[2] | $< \|L\| > = 0.48$, $< L^2 > = 0.30$ | Xtriage |
| Estimated twinning fraction | No twinning to report. | Xtriage |
| $F_o$,$F_c$ correlation | 0.97 | EDS |
| Total number of atoms | 2121 | wwPDB-VP |
| Average B, all atoms (Å$^2$) | 34.0 | wwPDB-VP |

Xtriage's analysis on translational NCS is as follows: *The largest off-origin peak in the Patterson function is 8.54% of the height of the origin peak. No significant pseudotranslation is detected.*

[1] Intensities estimated from amplitudes.

[2] Theoretical values of $< |L| >$, $< L^2 >$ for acentric reflections are 0.5, 0.333 respectively for untwinned datasets, and 0.375, 0.2 for perfectly twinned datasets.

For Manuscript Review

### 5 Model quality (i)

#### 5.1 Standard geometry (i)

Bond lengths and bond angles in the following residue types are not validated in this section: PEG, LAE

The Z score for a bond length (or angle) is the number of standard deviations the observed value is removed from the expected value. A bond length (or angle) with $|Z| > 5$ is considered an outlier worth inspection. RMSZ is the root-mean-square of all Z scores of the bond lengths (or angles).

| Mol | Chain | Bond lengths | | Bond angles | |
|---|---|---|---|---|---|
| | | RMSZ | #$\|Z\|$ >5 | RMSZ | #$\|Z\|$ >5 |
| 1 | A | 0.60 | 0/1954 | 0.96 | 0/2653 |

There are no bond length outliers.

There are no bond angle outliers.

There are no chirality outliers.

There are no planarity outliers.

#### 5.2 Too-close contacts (i)

In the following table, the Non-H and H(model) columns list the number of non-hydrogen atoms and hydrogen atoms in the chain respectively. The H(added) column lists the number of hydrogen atoms added and optimized by MolProbity. The Clashes column lists the number of clashes within the asymmetric unit, whereas Symm-Clashes lists symmetry-related clashes.

| Mol | Chain | Non-H | H(model) | H(added) | Clashes | Symm-Clashes |
|---|---|---|---|---|---|---|
| 1 | A | 1894 | 0 | 1875 | 9 | 0 |
| 2 | A | 17 | 0 | 19 | 0 | 0 |
| 3 | A | 14 | 0 | 20 | 3 | 0 |
| 4 | A | 196 | 0 | 0 | 1 | 0 |
| All | All | 2121 | 0 | 1914 | 9 | 0 |

The all-atom clashscore is defined as the number of clashes found per 1000 atoms (including hydrogen atoms). The all-atom clashscore for this structure is 2.

All (9) close contacts within the same asymmetric unit are listed below, sorted by their clash magnitude.

For Manuscript Review

| Atom-1 | Atom-2 | Interatomic distance (Å) | Clash overlap (Å) |
|---|---|---|---|
| 1:A:96:GLU:OE1 | 1:A:96:GLU:N | 2.25 | 0.65 |
| 1:A:194:LYS:NZ | 4:A:401:HOH:O | 2.28 | 0.62 |
| 1:A:194:LYS:CE | 3:A:302:PEG:H41 | 2.38 | 0.53 |
| 1:A:196:TYR:CG | 1:A:211:ARG:HD3 | 2.45 | 0.51 |
| 1:A:10:PHE:CD1 | 1:A:153:GLU:HB2 | 2.48 | 0.48 |
| 1:A:163:ARG:HG2 | 3:A:303:PEG:H21 | 1.99 | 0.44 |
| 1:A:196:TYR:CD1 | 1:A:211:ARG:HD3 | 2.54 | 0.43 |
| 1:A:202:ILE:HD11 | 3:A:302:PEG:H12 | 2.02 | 0.42 |
| 1:A:211:ARG:HD2 | 1:A:211:ARG:HA | 1.92 | 0.40 |

There are no symmetry-related clashes.

#### 5.3 Torsion angles (i)

##### 5.3.1 Protein backbone (i)

In the following table, the Percentiles column shows the percent Ramachandran outliers of the chain as a percentile score with respect to all X-ray entries followed by that with respect to entries of similar resolution.

The Analysed column shows the number of residues for which the backbone conformation was analysed, and the total number of residues.

| Mol | Chain | Analysed | Favoured | Allowed | Outliers | Percentiles |
|---|---|---|---|---|---|---|
| 1 | A | 236/241 (98%) | 236 (100%) | 0 | 0 | 100 100 |

There are no Ramachandran outliers to report.

##### 5.3.2 Protein sidechains (i)

In the following table, the Percentiles column shows the percent sidechain outliers of the chain as a percentile score with respect to all X-ray entries followed by that with respect to entries of similar resolution.

The Analysed column shows the number of residues for which the sidechain conformation was analysed, and the total number of residues.

| Mol | Chain | Analysed | Rotameric | Outliers | Percentiles |
|---|---|---|---|---|---|
| 1 | A | 199/201 (99%) | 199 (100%) | 0 | 100 100 |

There are no protein residues with a non-rotameric sidechain to report.

For Manuscript Review

Sometimes sidechains can be flipped to improve hydrogen bonding and reduce clashes. There are no such sidechains identified.

##### 5.3.3 RNA (i)

There are no RNA molecules in this entry.

#### 5.4 Non-standard residues in protein, DNA, RNA chains (i)

There are no non-standard protein/DNA/RNA residues in this entry.

#### 5.5 Carbohydrates (i)

There are no oligosaccharides in this entry.

#### 5.6 Ligand geometry (i)

3 ligands are modelled in this entry.

In the following table, the Counts columns list the number of bonds (or angles) for which Mogul statistics could be retrieved, the number of bonds (or angles) that are observed in the model and the number of bonds (or angles) that are defined in the Chemical Component Dictionary. The Link column lists molecule types, if any, to which the group is linked. The Z score for a bond length (or angle) is the number of standard deviations the observed value is removed from the expected value. A bond length (or angle) with $|Z| > 2$ is considered an outlier worth inspection. RMSZ is the root-mean-square of all Z scores of the bond lengths (or angles).

| Mol | Type | Chain | Res | Link | Bond lengths | | | Bond angles | | |
|---|---|---|---|---|---|---|---|---|---|---|
| | | | | | Counts | RMSZ | $\#\|Z\| > 2$ | Counts | RMSZ | $\#\|Z\| > 2$ |
| 2 | LAE | A | 301 | - | 17,17,17 | 0.44 | 0 | 17,21,21 | 0.52 | 0 |
| 3 | PEG | A | 303 | - | 6,6,6 | 0.24 | 0 | 5,5,5 | 0.36 | 0 |
| 3 | PEG | A | 302 | - | 6,6,6 | 0.31 | 0 | 5,5,5 | 0.42 | 0 |

In the following table, the Chirals column lists the number of chiral outliers, the number of chiral centers analysed, the number of these observed in the model and the number defined in the Chemical Component Dictionary. Similar counts are reported in the Torsion and Rings columns. '-' means no outliers of that kind were identified.

| Mol | Type | Chain | Res | Link | Chirals | Torsions | Rings |
|---|---|---|---|---|---|---|---|
| 2 | LAE | A | 301 | - | - | 0/13/23/23 | 0/1/1/1 |
| 3 | PEG | A | 303 | - | - | 2/4/4/4 | - |
| 3 | PEG | A | 302 | - | - | 1/4/4/4 | - |

For Manuscript Review

There are no bond length outliers.

There are no bond angle outliers.

There are no chirality outliers.

All (3) torsion outliers are listed below:

| Mol | Chain | Res | Type | Atoms |
|---|---|---|---|---|
| 3 | A | 303 | PEG | O1-C1-C2-O2 |
| 3 | A | 303 | PEG | C1-C2-O2-C3 |
| 3 | A | 302 | PEG | O2-C3-C4-O4 |

There are no ring outliers.

2 monomers are involved in 3 short contacts:

| Mol | Chain | Res | Type | Clashes | Symm-Clashes |
|---|---|---|---|---|---|
| 3 | A | 303 | PEG | 1 | 0 |
| 3 | A | 302 | PEG | 2 | 0 |

The following is a two-dimensional graphical depiction of Mogul quality analysis of bond lengths, bond angles, torsion angles, and ring geometry for all instances of the Ligand of Interest. In addition, ligands with molecular weight > 250 and outliers as shown on the validation Tables will also be included. For torsion angles, if less then 5% of the Mogul distribution of torsion angles is within 10 degrees of the torsion angle in question, then that torsion angle is considered an outlier. Any bond that is central to one or more torsion angles identified as an outlier by Mogul will be highlighted in the graph. For rings, the root-mean-square deviation (RMSD) between the ring in question and similar rings identified by Mogul is calculated over all ring torsion angles. If the average RMSD is greater than 60 degrees and the minimal RMSD between the ring in question and any Mogul-identified rings is also greater than 60 degrees, then that ring is considered an outlier. The outliers are highlighted in purple. The color gray indicates Mogul did not find sufficient equivalents in the CSD to analyse the geometry.

For Manuscript Review

Ligand LAE A 301

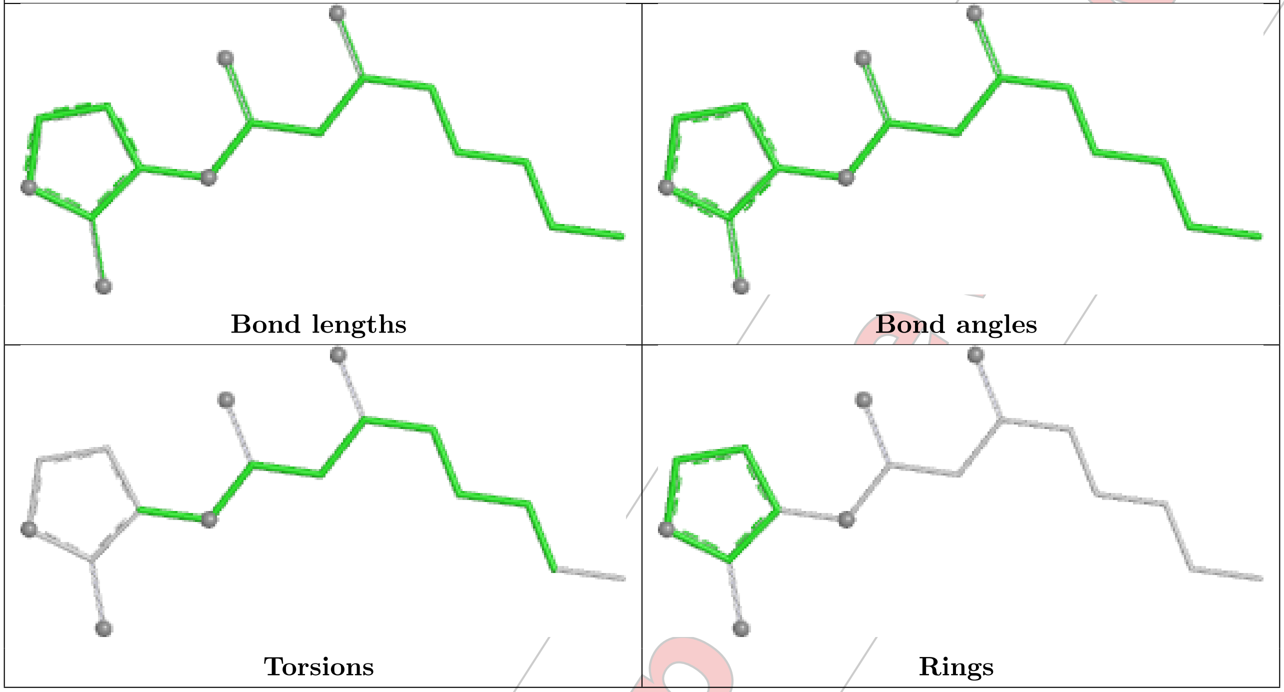


#### 5.7 Other polymers (i)

There are no such residues in this entry.

#### 5.8 Polymer linkage issues (i)

There are no chain breaks in this entry.

For Manuscript Review

### 6 Fit of model and data (i)

#### 6.1 Protein, DNA and RNA chains (i)

In the following table, the column labelled '#RSRZ> 2' contains the number (and percentage) of RSRZ outliers, followed by percent RSRZ outliers for the chain as percentile scores relative to all X-ray entries and entries of similar resolution. The OWAB column contains the minimum, median, $95^{th}$ percentile and maximum values of the occupancy-weighted average B-factor per residue. The column labelled 'Q< 0.9' lists the number of (and percentage) of residues with an average occupancy less than 0.9.

| Mol | Chain | Analysed | <RSRZ> | #RSRZ>2 | | | OWAB(Å$^2$) | Q<0.9 |
|---|---|---|---|---|---|---|---|---|
| 1 | A | 233/241 (96%) | 0.58 | 14 (6%) | 29 | 28 | 14, 31, 48, 60 | 7 (3%) |

All (14) RSRZ outliers are listed below:

| Mol | Chain | Res | Type | RSRZ |
|---|---|---|---|---|
| 1 | A | 2 | ILE | 8.2 |
| 1 | A | 97 | PRO | 3.3 |
| 1 | A | 219 | PHE | 2.8 |
| 1 | A | 95 | VAL | 2.6 |
| 1 | A | 178 | ALA | 2.5 |
| 1 | A | 1 | MET | 2.4 |
| 1 | A | 177 | PRO | 2.4 |
| 1 | A | 74 | VAL | 2.4 |
| 1 | A | 45 | GLN | 2.4 |
| 1 | A | 114 | GLY | 2.3 |
| 1 | A | 240 | ASN | 2.3 |
| 1 | A | 3 | GLU | 2.2 |
| 1 | A | 211 | ARG | 2.1 |
| 1 | A | 136 | ALA | 2.1 |

#### 6.2 Non-standard residues in protein, DNA, RNA chains (i)

There are no non-standard protein/DNA/RNA residues in this entry.

#### 6.3 Carbohydrates (i)

There are no oligosaccharides in this entry.

For Manuscript Review

#### 6.4 Ligands (i)

In the following table, the Atoms column lists the number of modelled atoms in the group and the number defined in the chemical component dictionary. The B-factors column lists the minimum, median, $95^{th}$ percentile and maximum values of B factors of atoms in the group. The column labelled 'Q< 0.9' lists the number of atoms with occupancy less than 0.9.

| Mol | Type | Chain | Res | Atoms | RSCC | RSR | B-factors($Å^2$) | Q<0.9 |
|---|---|---|---|---|---|---|---|---|
| 3 | PEG | A | 303 | 7/7 | 0.83 | 0.14 | 39,48,53,54 | 0 |
| 3 | PEG | A | 302 | 7/7 | 0.85 | 0.12 | 42,46,53,54 | 0 |
| 2 | LAE | A | 301 | 17/17 | 0.95 | 0.08 | 24,26,36,38 | 0 |

The following is a graphical depiction of the model fit to experimental electron density of all instances of the Ligand of Interest. In addition, ligands with molecular weight > 250 and outliers as shown on the geometry validation Tables will also be included. Each fit is shown from different orientation to approximate a three-dimensional view.

**Electron density around LAE A 301:**

$2mF_o$-$DF_c$ (at 0.7 rmsd) in gray
$mF_o$-$DF_c$ (at 3 rmsd) in purple (negative) and green (positive)

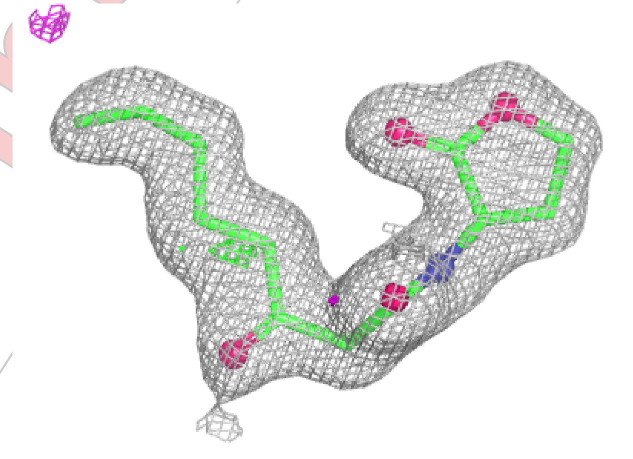

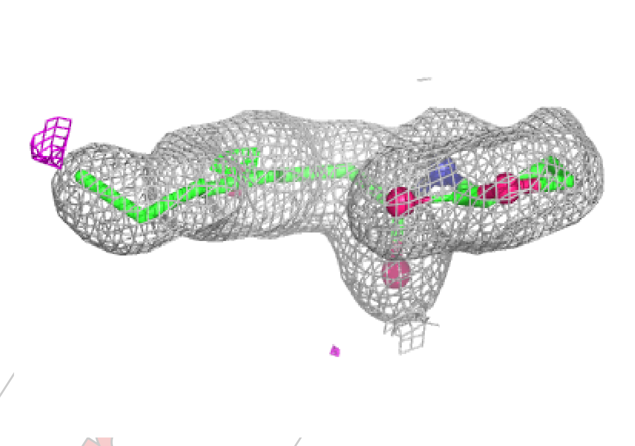

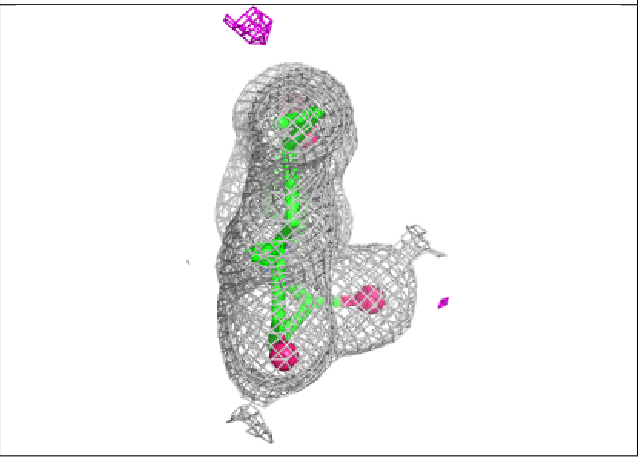

#### 6.5 Other polymers (i)

There are no such residues in this entry.

For Manuscript Review
